## Supplementary Information for "Droplet growth, Ostwald’s rule, and emergence of order in Fused in Sarcoma"

#### Supplementary Methods

**SOP-IDP model:** The simulations of the FUS proteins were carried out using the transferable Self-Organized Polymer (SOP-IDP) model [1], which is an improved version of the model [2]. In the SOP-IDP model, each residue is represented using two interacting sites: a backbone bead at the center of the $C_\alpha$ atom and a side-chain bead located on the center of mass of the side-chain. The SOP-IDP energy function is given by,

$$\begin{aligned} U_{SOP-IDP} &= U_{FENE} + U_{ELE} + U_{EXV} + U_{BB} + U_{BS} + U_{SS} \\ &= -\sum_{i=1}^{N_B} \frac{k}{2} R_0^2 \log\left(1 - \frac{(r_i - r_{ref,i})^2}{R_0^2}\right) + \sum_{i,j} \frac{e_i e_j exp(-\kappa r_{ij})}{\epsilon r_{ij}} + \sum_{i=1}^{N_{loc}} \varepsilon_{loc}\left(\frac{\sigma_i}{r_i}\right)^6 \\ &+ \sum_{i=1}^{N_{BB}} \varepsilon_{BB}\left[\left(\frac{\sigma_i}{r_i}\right)^{12} - 2\left(\frac{\sigma_i}{r_i}\right)^6\right] + \sum_{i=1}^{N_{BS}} \varepsilon_{BS}\left[\left(\frac{\sigma_i}{r_i}\right)^{12} - 2\left(\frac{\sigma_i}{r_i}\right)^6\right] \\ &+ \sum_{i=1}^{N_{SS}} \varepsilon_{SS}|\varepsilon_i - 0.7|\left[\left(\frac{\sigma_i}{r_i}\right)^{12} - 2\left(\frac{\sigma_i}{r_i}\right)^6\right]. \end{aligned} \quad (1)$$

The first term, $U_{FENE}$, in Eq. 1, with $r_{ref,i}$ denoting the equilibrium distance between bonded moieties (such as adjacent backbone beads and side-chains attached to the backbone beads), represents the finitely extensible nonlinear elastic (FENE) potential, which maintains chain connectivity. We set $k = 10$ kcal/mol/Å$^2$ is the bond stiffness, $R_0 = 2$Å is the tolerance for fluctuations around the equilibrium bond length, and $N_B$ is the total number of covalent bonds in a polypeptide chain. The second term, $U_{ELE}$, accounting for the electro-

Table S1: **Radii and charges for backbone and side-chain beads.** Parameters for the coarse-grained beads in the SOP-IDP model.

| Residue | Radius (Å) | Charge (e) |
|---|---|---|
| Ala | 2.52 | 0.0 |
| Val | 2.93 | 0.0 |
| Leu | 3.09 | 0.0 |
| Ile | 3.09 | 0.0 |
| Met | 3.09 | 0.0 |
| Phe | 3.18 | 0.0 |
| Pro | 2.78 | 0.0 |
| Ser | 2.59 | 0.0 |
| Thr | 2.81 | 0.0 |
| Asn | 2.84 | 0.0 |
| Gln | 3.01 | 0.0 |
| Tyr | 3.23 | 0.0 |
| Trp | 3.39 | 0.0 |
| Asp | 2.79 | -1.0 |
| Glu | 2.96 | -1.0 |
| His | 3.04 | 0.0 |
| Lys | 3.18 | 1.0 |
| Arg | 3.28 | 1.0 |
| Cys | 2.74 | 0.0 |
| Gly | 0.00 | 0.0 |
| Backbone | 1.90 | 0.0 |

static interactions between all pairs of charged residues, is modeled using the Debye-Hückel potential. The point charges $e_i$ are located at the center of the side chain bead of the charged residues (see Table S1). The inverse Debye length, $\kappa$, is calculated at 150 mM NaCl salt concentration (used in the simulations) and the dielectric constant $\epsilon$ was taken to be 78. The third term in Eq. 1 represents the repulsive excluded volume interactions that prevents overlap of beads. The final three terms in the energy function capture the sequence-specific interactions, detailing backbone-backbone (BB), backbone-side-chain (BS), and side-chain-side-chain (SS) interactions, respectively. The precise sequence is encoded by the chemical identity of the side chains. The parameter $\varepsilon_i$ in Eq. 1 is based on the statistical potential [3]. The three parameters in the SOP-IDP model, $\varepsilon_{BB}$, $\varepsilon_{BS}$ and $\varepsilon_{SS}$ were determined using a learning procedure [1].

**Monomer simulations:** We performed all the simulations using the SOP-IDP model [1]. The equilibrium ensembles of the monomers were generated using the low friction Langevin dynamics simulations as implemented in the LAMMPS package [4]. The equation of motion, for bead $i$ at position $r_i$ is,

$$m_i \ddot{r}_i = -\frac{\partial U_{SOP-IDP}}{\partial r_i} - \gamma m_i \dot{r}_i + \Gamma_i \tag{2}$$

where $\gamma$ is the friction coefficient, $\Gamma$ is a Gaussian random force which satisfies the fluctuation-dissipation relation, $< \Gamma_i(t)\Gamma_j(t') > = 6k_BT\gamma\delta_{ij}\delta(t-t')$, $m_i$ is the mass of the bead. We integrated Eq. 2 using the velocity-Verlet algorithm [5], as implemented within the LAMMPS package, using a time-step of 10 fs. To enhance the conformational sampling [6], the friction coefficient $\gamma$ was set to 0.5 $ps^{-1}$, which is around 1% of the friction in bulk water. The monomer simulations were carried out for a total of $2 \times 10^9$ steps. The first $3 \times 10^8$ steps were considered as equilibration time. We collected statistics every 10,000 steps for the rest of the trajectory. All monomer simulations for monomers were carried out at 150 mM NaCl and at room temperature.

**Identification of $N^*$ states:** Conformations with propensity to form fibrils, that belong to $N^*$ states, were identified using the equilibrium monomer conformational ensembles by comparing their structural overlap (Eq. 3) with the monomer units of the experimentally determined structures. The S-bend topology was used for core-1 (residues 39-95; PDB ID: 5W3N) and the U-bend topology for core-2 (residues 112-150; PDB ID: 6XFM). For core-3, we used the predicted structure in this work (residues 155-190; see the Main text). We computed the structural overlap $\chi_{fib}$ at time step $\tau$ using,

$$\chi_{fib}(\tau) = \frac{1}{N_{pairs}} \sum_{a,b}^{N_{pairs}} \Theta(d - |r_{a,b}(\tau) - r^0_{a,b}|), \tag{3}$$

where $r^0_{a,b}$ is the distance between sites $a$ and $b$ in the reference (experimentally determined or predicted) structure, $\Theta$ is the Heaviside step function. Following our previous studies [7, 8], we identified conformations with $\chi_{fib}(\tau) \geq \chi_c$ as having the propensity to adopt fibril-like structures and are classified as the $N^*$ state.

We chose the cutoff values $\chi_{c.i}$ based on the following rationale. This threshold defines the onset of fibril-like conformations for which $\chi_{fib.i} \geq \chi_{c.i}$. Such conformations are likely to be similar to the reference fibril structure. If this holds good we expect the mean first passage time $\langle \tau_{c.i} \rangle$ should remain constant for all $\chi_{fib.i} \geq \chi_{c.i}$, since these conformations belong to the same energy minimum. Therefore, for each core $i$, we selected $\chi_{c.i}$ such that $\langle \tau_{c.i} \rangle$ reaches a plateau and does not change with upon further increases in $\chi_{c.i}$ (see Fig. S1A).

**Relative stabilities of core-1, core-2, and core-3.** The stability of a core structure was also determined using the overlap function $\chi_{fib}$ (see Eq. 3). Using all the fibril-like conformations (microstates) of core $i$ (macrostate), where $i = 1, 2, 3U, 3S$, generated in the Langevin simulations of the full-length FUS-LC, we calculated the probability of forming a fibril-like structure (a single macrostate), using the method used previously [9],

$$P_{c.i} = \frac{\sum_m \Theta(\chi_m - \chi_{c.i}) e^{-\beta U_m}}{\sum_m e^{-\beta U_m}}, \tag{4}$$

where $\beta = 1/k_B T$. The sum in Eq.4 is over all the $m$ conformations that are generated in the simulations. The Heaviside step function $\Theta$ enforces a zero contribution from non-fibril-like conformations in the numerator. Using the calculated probabilities, the relative free energy change between two cores $i$ and $j$ is,

$$\Delta F_{i,j} = -k_B T \ln \left( \frac{P_{c.i}}{P_{c.j}} \right). \tag{5}$$

**Contact Maps:** Residues $i$ and $j$ are considered to be in contact if the distance between their respective side-chains is $\leq$ 0.8 nm. The time-dependent inter-residue matrices are transformed into probability contact maps $p_{ij}(\tau)$ using the logistic function,

$$p_{ij}(\tau) = \frac{1}{1 + exp[\beta(r_{ij}(\tau) - r_0)]} \tag{6}$$

where $\beta$ = 500 nm$^{-1}$, and $r_0$ = 0.8 nm. (The actual value of $\beta$ is unimportant as long as it is large.) We obtained the ensemble-averaged probability map, $\langle p_{ij}^X \rangle$, for the IDP $X$ ($X$ is either FUS-LC, or FUS-LC-N, or CTC) by averaging over all the snapshots in all trajectories.

**Local free energy minima:** The conformations obtained from the Brownian Dynamics (BD) trajectories were divided into distinct microstates through a structural clustering algorithm. Initially, each conformation was converted to the corresponding distribution of reciprocal interatomic distance (DRID) metric [10], which maintains the kinetic distance between any two conformations. After mapping into the DRID space, the conformational ensemble was clustered using a regular space clustering algorithm implemented in the PyEMMA 2.5 distribution [11]. A cutoff value of 1.1 was applied to identify the distinct microstates.
The free energy of minimum $i$ was estimated as $\Phi_i = -k_B T ln[Z_i]$, where $Z_i$ is the partition function, defined as the number of conformations $N_i$ in cluster $i$ [12]. The complete partition function, $Z$, corresponds to the total count of conformations, $N$, generated in the BD trajectories.

**Kinetic rate constants:** To estimate the relative reverse rate constants from fibril-like states to the disordered ensemble (random coil, RC), we used the mean first passage times (MFPTs) from the RC ensemble to each fibril conformation ($\tau_i$) and the free energy differences $\Delta G_{i3} = G_i - G_3$ (where $i$=1,2) relative to the S-bend core-3. Assuming detailed balance and a two-state transition pathway between the RC and each fibril conformation, the forward and reverse rate constants are related by

$$\frac{k_{\text{fwd},i}}{k_{\text{rev},i}} = e^{-\beta \Delta G_i}, \tag{7}$$

where $\Delta G_i$ is the free energy difference between state $i$ and the RC ensemble. We do not need to know $\Delta G_i$ directly if we only require the relative reverse rate constants. Given that $k_{\text{fwd},i} = \frac{1}{\tau_i}$ and choosing core-3 as the reference, the ratio of reverse rate constants becomes

$$\frac{k_{\text{rev},i}}{k_{\text{rev},3}} = \frac{\tau_3}{\tau_i} \cdot e^{\Delta G_{i3}/RT}, \tag{8}$$

where $\Delta G_{i3} = G_i - G_3$, and $\tau_i$ is the MFPT from RC to fibril state $i$.

The final computed ratios of reverse rate constants, along with uncertainties propagated from the standard errors of MFPTs and free energy differences, are summarized in the table S2.

**Concentrations of the two phases.** To estimate the volume occupied by droplets in each simulation frame, we employed a grid-based method inspired by phase imaging techniques. In Quantitative Phase Imaging (QPI), variations in optical path length—proportional to the local refractive index and, by extension, to the local mass density—are used to reconstruct a refractive index map. This enables the identification and segmentation of dense regions such as intracellular condensates or protein droplets. Analogously, we discretized the simulation box into a three-dimensional Cartesian grid with uniform voxel size $v = l^3$, where $l = 9.2$, Å is the voxel edge length. Each droplet was defined by the positions of all its constituent beads. Let $N_v$ be the number of voxels occupied by at least one bead. The total volume $V$ of the droplet is then given by:

$$V = N_v \cdot v \tag{9}$$

This approach provides a resolution-controlled estimate of the spatial extent of droplets, conceptually similar to how QPI identifies dense material regions based on local optical phase shifts. To estimate the mass concentration within each droplet, we computed the total mass $M$ as:

$$M = N_c \cdot m_c \tag{10}$$

where $N_c$ is the number of chains in the droplet and $m_c$ is the mass of a single chain. The molar concentration $C$ inside the droplet is then calculated as:

$$C = \frac{M}{N_A \cdot V} \tag{11}$$

where $N_A$ is Avogadro's number. The resulting concentration was reported in millimolar (mM) after appropriate unit conversion.

**Chemical potentials of the two phases:** A minimal criterion to establish the coexistence between two phases is to ensure that their chemical potentials are the same. To compute the chemical potential ($\mu$) in both the condensed and dilute phases, we used a modified version of the Widom particle insertion approach [13]. In the calculation of $\mu$, we do not directly insert proteins into the droplet, as done in the traditional Widom method, for several reasons:

1. The effect of such insertion would depend on the conformation of the intrinsically disordered protein (IDP) monomer.

2. The specific residues of the inserted chain conformation would interact differently with the rest of the droplet.

3. Insertion of polymers could result in steric repulsion with other monomers in the droplet, requiring equilibration at each step.

Because it is generally recommended to equilibrate the system after insertion before measuring its free energy [14], this approach presents challenges in our two-phase coexistence simulations. The continuous exchange of protein chains between the dilute and condensed phases means that prolonged equilibration might alter the system by changing the number of chains in the droplet by more or less than the required by the Widom method. Instead, we used the conformations sampled in our simulations, which is justified because all the oligomers and droplets formed stochastically without any bias, allowing us to consider these conformations as equilibrated. Furthermore, a simple average of the chemical potentials computed using the particle insertion [13] and particle deletion [15] method may not be accurate for the the system studied here [16]. It is suggested that only the insertion scheme should be considered for accurate results [17, 18]. To ensure the accuracy of our calculations, we consider only those consecutive snapshots $\tau, \tau+1$ where the number of chains in the phase $p$ (condensed or dilute) increased by 1, from $N$ at $\tau$ to $N+1$ at $\tau+1$. This approach mimics the particle insertion scheme, although in our case it represents protein "insertion".

The chemical potential is computed for the largest oligomer (or droplet) and for the dilute phase at time step $\tau$ as:

$$\mu_p = \left(\frac{\partial F_p}{\partial N_p}\right)_{T,V}. \quad (12)$$

The change in free energy associated with the insertion of a protein into the phase $p$ is given by $\Delta F_p = -k_B T ln \left\langle e^{-\beta \Delta U_p} \right\rangle$ [5], where $\Delta U_p = U_p(\tau+1, N_p+1) - U_p(\tau, N_p)$ ($\equiv \Delta U_{SOP-IDP.p}$) is the change in the potential energy, $k_B$ is the Boltzmann constant. The modified approach allows us to calculate the chemical potential in both the phases while overcoming the unique challenges in the simulations of multichain IDP systems, while maintaining the theoretical basis of the particle insertion method.

**Additional Analysis:** The bin sizes for all the histograms were determined based on the Freedman-Diaconis rule [19] to ensure optimal representation of the data. For nonparametric density estimation, a normalized Gaussian kernel function was employed, with the bandwidth set to $h = N^{-1/5}$, as recommended elsewhere [20]. The secondary structures were assigned based on the positions of the $C_\alpha$ atoms using the protein C-alpha secondary structure output (PCASSO) algorithm [21]. Throughout the text, unless otherwise specified, error bars represent the standard error of the mean.

*Nematic Order Parameter:* To assess if there is orientational order in the droplet phase, we computed the nematic order parameter using,

$$S_2 = \frac{1}{2}\left\langle 3\cos^2\theta - 1\right\rangle, \tag{13}$$

where $\theta$ is the angle between a molecule's long axis and the director (the preferred orientation direction in the nematic phase), and the angle brackets denote an ensemble average. As defined in the above equation, $S_2$ satisfies, $-0.5 \leq S_2 \leq 1$. $S_2$ = -0.5 represents perfect perpendicular alignment, $S_2 = 0.0$ indicates a completely random orientation (isotropic phase), and $S_2 = 1.0$ corresponds to perfect parallel alignment.

*Jensen-Shannon divergence:* The JSD, ($D_{JS}$), a measure of similarity between two distribution functions, $P_d(\mu)$ and $P_c(\mu)$, and is calculated using,

$$D_{JS}(P_d(\mu) \parallel P_c(\mu)) = \frac{1}{2}D_{KL}(P_d(\mu) \parallel P(\mu)) + \frac{1}{2}D_{KL}(P_c(\mu) \parallel P(\mu)), \tag{14}$$

where $P(\mu) = \frac{1}{2}\big(P_d(\mu) + P_c(\mu)\big)$ is the pointwise average of the two distributions. The Kullback-Leibler divergence($D_{KL}$) is defined as $D_{KL}(P_d(\mu) \parallel P_c(\mu)) = \sum_\mu P_d(\mu)\log\frac{P_d(\mu)}{P_c(\mu)}$.

*Coefficient of variation:* The squared coefficient of variation of $\mu$, a measure of fluctuations, is

$$\sigma_\mu = (\langle\mu^2\rangle - \langle\mu\rangle^2)/\langle\mu\rangle^2 \tag{15}$$

**Solvent-accessible surface area (SASA):** We calculated SASA computed using the Shrake-Rupley algorithm as implemented in MDTraj [22]. A probe radius of 0.14 nm and 100 surface points per atom were used. For $SASA$ calculations, custom atomic radii corresponding to the coarse-grained beads in the SOP-IDP model (see Table S1) were supplied to MDTraj to ensure accurate surface estimation. The total $SASA$ of each monomer was obtained by summing the contributions from all the beads.

*Gyration tensor and shape parameter:* We calculated the gyration tensor for the entire droplet. Using the eigenvalues $\lambda_g^2$ $(g = 1, 2, 3)$ of the gyration tensor $S_{xy}$,

$$S_{xy}^{(j)} = \frac{1}{2N_j^2} \sum_i \sum_k (x_i - x_k)(y_i - y_k), \tag{16}$$

To characterize the shape of a droplets, we calculated the relative shape anisotropy

$$\kappa^2 = \frac{3}{2} \cdot \frac{\lambda_1^4 + \lambda_2^4 + \lambda_3^4}{(\lambda_1^2 + \lambda_2^2 + \lambda_3^2)^2} - \frac{1}{2} \tag{17}$$

$\kappa^2$ ranges from 0 to 1, where $\kappa^2 = 0$ corresponds to a perfectly spherical shape, and $\kappa^2 = 1$ indicates that all points lie along a straight line.

### $N^*$ theory

The $N^*$ theory posits that within the native basin of attraction of the monomer, sparsely populated higher free-energy conformations—excited states relative to the lowest-energy minimum exist. There are collectively referred to as $N^*$ states, which resemble structural motifs found in amyloid fibrils [23, 24, 25]. These transient excitations in the free energy landscape act as assembly-competent precursors to aggregation [24]. Although typically accounting for less than 1% of the monomer ensemble under physiological conditions, $N^*$ states play a central role in initiating the aggregation cascade by serving as structural templates for oligomer formation [26, 27]. In this framework, fibril formation is governed by the kinetics of accessing and stabilizing these $N^*$ conformations. The time scale for aggregation depends exponentially on the population of such fibril-like states [24]. Thus, the $N^*$ concept provides a unifying explanation for fibril polymorphism and sequence-dependent aggregation propensities [7].

### Supplementary Results

#### Properties of FUS-LC condensates

**FUS-LC chains in condensates are dynamic:** The dynamic nature of droplets is highlighted by the absence of persistent internal order, as measured by the nematic order parameter $S_2$ (Eq. 13). Small oligomers, particularly dimers and trimers, exhibit a modest degree of orientational order with $S_2$ values of 0.25 (Fig. S12). This increased order is likely driven by the necessity for internal FUS-LC molecules to arrange themselves in a manner that optimizes inter-chain interactions, which occurs more readily when the chains are aligned. However, the orientational preference comes at the cost of reduced entropy. As the number of chains in the oligomers increase, $S_2$ decreases, eventually becoming $\approx 0$ in the droplet. This finding is rationalized by noting that in the larger assemblies, internal FUS-LC molecules can form interactions with numerous other chains without reduction in orientational entropy. The average value of $S_2 = 0$ for monomers inside the droplets (inset in Fig. S12) implying that molecular orientations within the simulated condensates are essentially random. The lack of long-range orientational order in FUS-LC condensates is consistent with the notion that intrinsically disordered proteins (IDPs) are flexible and are devoid of stable tertiary structure. The absence of orientational order with liquid-like properties of biomolecular condensates of IDPs in which polymer chains diffuse freely within the droplet.

#### Structural models of FUS-LC fibrils predicted by AlphaFold

We performed structural predictions on multidomain complexes with varying numbers of chains: $FUS_{39\text{-}95}$ and $FUS_{1\text{-}110}$ for core-1, and $FUS_{112\text{-}150}$ and $FUS_{111\text{-}214}$ for core-2. Each chain count was assessed using eight distinct random seeds. AlphaFold provides quality metrics—pTM, iPTM, and pLDDT [28]—which exhibited substantial variability across different chain conformations. To identify the optimal structural predictions for core-1 and core-2, we examined how the distributions of these metrics changed with chain number, thereby evaluating prediction reliability and confidence (see Fig. S13).

**Comparison of structural model of core-1:**. The structures generated by AlphaFold3

(AF3) are illustrated in Figure 5B in the main text, where the fibril core is formed from the W-shaped monomers. In this model, the outer core segments correspond to residues 39–47 and 83–95, while the inner core segments are defined by residues 53–63 and 67–78. The AF3-derived structural model identifies residues 39–47, 53–63, 67–70, 76-79 and 83–95 as forming the $\beta$-strands of a parallel superpleated $\beta$-structure. The experimental structure ([29]; Fig. 5A) reveals extended outer segments comprising residues 39–55 and 84–95, with a compact inner core spanning residues 62–78. Residues 39–41, 52–55, 62–64, 66–70, 85–90, and 92–95 were identified as forming the $\beta$-strands with serpentine-like $\beta$-sheets. We assigned $\beta$-strand using backbone dihedral angles and hydrogen-bond patterns, analyzed using the Visual Molecular Dynamics (VMD) software [30]. In the AF3-predicted structure, residues 48–52, 64-67, and 79-82 adopt a $\beta$-turn conformations, facilitating chain reversal, while residues 70-76 form irregular region. The structural overlap computed for residues 39-95 from AF3 prediction with respect to ssNMR structure is $\chi = 0.19$ and RMSD value for all $C_\alpha = 13.8$Å. We also used AF3 to predict the structure of residues 39–95 (Fig. S15C), corresponding to core-1. This resulted in poorer agreement with the experimental structure, yielding $\chi = 0.15$ and RMSD values for all $C_\alpha$ atoms of 16.9Å.

Although the structural prediction from AF3 (Fig. 5B) deviates from the experimental ssNMR structure (PDB ID: 5W3N), it is sufficiently accurate to validate the Ostwald's rule of stages. We arrive at this conclusion by comparing the MFPTs using the AF3-predicted and experimental structures. The raw MFPT values show remarkable agreement: $\langle\tau_{c.1}^{\mathrm{AF3}}\rangle = 121.42 \pm 8.67$ ms vs. $\langle\tau_{c.1}^{\mathrm{exp}}\rangle = 129.17 \pm 11.73$ ms. Application of the correction formula (Eq. 2) further reinforces this correspondence: $\langle\widehat{\tau_{c.1}^{\mathrm{AF3}}}\rangle = 1974.97 \pm 13.36$ ms (corrected AF3) vs. $\langle\widehat{\tau_{c.1}^{\mathrm{exp}}}\rangle = 2134.0 \pm 16.1$ ms. Although the values are not identical, both are sufficiently large to support the conclusion that core-1 forms much slower than core-2 and core-3.

**Assessing the accuracy AlphaFold core-2 structures:** In the structures generated by AF3 for $\mathrm{FUS}_{111\text{-}214}$ and $\mathrm{FUS}_{112\text{-}150}$, the outer and inner extended segments correspond to residues 111–131 and 135–150, respectively (Figs. S15B and D). Similarly, the experimental structure defines these segments as residues 112–127 (outer) and 132–150 (inner) (Fig. S15A). The AF3-derived computational model identifies residues 115–123, 129–131, 135–136, and 139–148 as forming the $\beta$-strands of parallel $\beta$-sheets. These are in reasonable agreement with the AlphaFold3 (AF3; [28]) experimental data, where residues 113–122, 135–136, and 139–149 are predicted to be $\beta$-strands. In the AF3-predicted structure, residues 132–134 adopt a $\beta$-turn conformation, facilitating chain reversal within the irregular region (residues 123–134). Glycines at positions 137 and 138 (G137/G138) partition the 135–148 segment into two discontinuous $\beta$-strands in both computational and experimental models, even though the entire stretch maintains a fully extended backbone conformation. The structural overlap for residues 112-150, computed from the AF3 prediction of FUS 111-214 relative to the experimental structure, is $\chi = 0.26$, with an RMSD of 8.9Å for all $C_\alpha$ atoms. We also used AF3 to predict the structure of just the residues 112-150 (Fig. S15D), corresponding to core-2. This resulted in similar agreement with the experimental structure, yielding $\chi = 0.29$ and RMSD values for all $C_\alpha$ atoms of 9.3Å.

To ascertain that validity of Ostwald's rule of stages, we calculated the MFPTs using the predicted structures for all the cores. The values of the MFPTs obtained from the trajectories that the reached the three cores are: $\langle \tau^{\mathrm{AF3}}_{c.2|s} \rangle = 104.98 \pm 4.56$ ms vs. $\langle \tau^{\mathrm{exp}}_{c.2|s} \rangle = 97.47 \pm 8.70$ ms. By using the correction formula (Eq. 2) demonstrates the disagreement: $\langle \widehat{\tau^{\mathrm{AF3}}_{c.2}} \rangle = 377.59 \pm 8.73$ ms (corrected AF3) vs. $\langle \widehat{\tau^{\mathrm{exp}}_{c.2}} \rangle = 929.0 \pm 9.8$ ms.

#### Structural models of FUS-LC fibrils predicted by AlphaFold2

To investigate structural variations, we predicted $\mathrm{FUS}_{155\text{-}190}$ and $\mathrm{FUS}_{141\text{-}214}$ fibrils using AF2 [31] by applying the same analysis as described above (see Fig. S17). While the AF2 predictions maintained a U-bend topology within the monomers, the fibril structure deviated from the canonical architecture observed in core-1 and core-2 fibrils [29, 32]. In particular, it lacks the typical parallel alignment of protein subunits (see Figs. S19A and S20A). Nonetheless, AF2 models consistently displayed the U-bend topology, with a conserved "kinked" region spanning residues 176-184. The maximum values for iPTM, pTM, and average pLDDT across all AF2 predictions were notably lower than those obtained from AF3 predictions, with values of 0.16, 0.18, and 23.90, respectively (see Table **??**). Despite these differences, the intra- and inter-chain contact maps were similar to those predicted by AF3, with residues 164-176 showing both a higher probability and a longer range of intermolecular contacts than other residues (see Figs. S19B and S20B). All models exhibited an overall lower $\beta$-strand propensity compared to AF3 models, though the propensity remained relatively high (see Figs. S19C and S20C). This consistency reinforces the prominence of the U-bend motif across different predictive frameworks.

### References

[1] Mauro L Mugnai et al. "Sizes, conformational fluctuations, and SAXS profiles for Intrinsically Disordered Proteins". In: *Prot. Sci.* 34 (2025), e70067.

[2] Upayan Baul et al. "Sequence effects on size, shape, and structural heterogeneity in intrinsically disordered proteins". In: *The Journal of Physical Chemistry B* 123.16 (2019), pp. 3462–3474.

[3] Marcos R Betancourt and D Thirumalai. "Pair potentials for protein folding: choice of reference states and sensitivity of predicted native states to variations in the interaction schemes". In: *Protein science* 8.2 (1999), pp. 361–369.

[4] Steve Plimpton. "Fast parallel algorithms for short-range molecular dynamics". In: *Journal of computational physics* 117.1 (1995), pp. 1–19.

[5] Daan Frenkel and Berend Smit. *Understanding molecular simulation: from algorithms to applications*. Elsevier, 2023.

[6] JD Honeycutt and D Thirumalai. "The nature of folded states of globular proteins". In: *Biopolymers: Original Research on Biomolecules* 32.6 (1992), pp. 695–709.

[7] Abhinaw Kumar et al. "Sequence determines the switch in the fibril forming regions in the low-complexity FUS protein and its variants". In: *The journal of physical chemistry letters* 12.37 (2021), pp. 9026–9032.

[8] Debayan Chakraborty, John E Straub, and D Thirumalai. "Energy landscapes of A$\beta$ monomers are sculpted in accordance with Ostwald's rule of stages". In: *Science Advances* 9.12 (2023), eadd6921.

[9] DK Klimov and D Thirumalai. "Cooperativity in protein folding: from lattice models with sidechains to real proteins". In: *Folding and Design* 3.2 (1998), pp. 127–139.

[10] Ting Zhou and Amedeo Caflisch. "Distribution of reciprocal of interatomic distances: A fast structural metric". In: *Journal of Chemical Theory and Computation* 8.8 (2012), pp. 2930–2937.

[11] Martin K Scherer et al. "PyEMMA 2: A software package for estimation, validation, and analysis of Markov models". In: *Journal of chemical theory and computation* 11.11 (2015), pp. 5525–5542.

[12] Sergei V Krivov and Martin Karplus. "Free energy disconnectivity graphs: Application to peptide models". In: *The Journal of chemical physics* 117.23 (2002), pp. 10894–10903.

[13] Ben Widom. "Some topics in the theory of fluids". In: *The Journal of Chemical Physics* 39.11 (1963), pp. 2808–2812.

[14] Georgios C Boulougouris, Ioannis G Economou, and Doros N Theodorou. "On the calculation of the chemical potential using the particle deletion scheme". In: *Molecular physics* 96.6 (1999), pp. 905–913.

[15] KS Shing and KE Gubbins. "The chemical potential in dense fluids and fluid mixtures via computer simulation". In: *Molecular Physics* 28.5 (1892), pp. 1109–1128.

[16] Nandou Lu and David A Kofke. "Accuracy of free-energy perturbation calculations in molecular simulation. I. Modeling". In: *The Journal of Chemical Physics* 114.17 (2001), pp. 7303–7311.

[17] Nandou Lu and David A Kofke. "Accuracy of free-energy perturbation calculations in molecular simulation. II. Heuristics". In: *The Journal of Chemical Physics* 115.15 (2001), pp. 6866–6875.

[18] Nandou Lu, Jayant K Singh, and David A Kofke. "Appropriate methods to combine forward and reverse free-energy perturbation averages". In: *The Journal of Chemical Physics* 118.7 (2003), pp. 2977–2984.

[19] David Freedman and Persi Diaconis. "On the histogram as a density estimator: L 2 theory". In: *Zeitschrift für Wahrscheinlichkeitstheorie und verwandte Gebiete* 57.4 (1981), pp. 453–476.

[20] David W Scott. *Multivariate density estimation: theory, practice, and visualization.* John Wiley & Sons, 2015.

[21] Sean M Law, Aaron T Frank, and Charles L Brooks III. “PCASSO: A fast and efficient Cα-based method for accurately assigning protein secondary structure elements”. In: *Journal of computational chemistry* 35.24 (2014), pp. 1757–1761.

[22] Robert T. McGibbon et al. “MDTraj: A Modern Open Library for the Analysis of Molecular Dynamics Trajectories”. In: *Biophysical Journal* 109.8 (2015), pp. 1528–1532. DOI: 10.1016/j.bpj.2015.08.015.

[23] John E Straub and D Thirumalai. “Toward a molecular theory of early and late events in monomer to amyloid fibril formation”. In: *Annual review of physical chemistry* 62.1 (2011), pp. 437–463.

[24] Mai Suan Li et al. “Factors governing fibrillogenesis of polypeptide chains revealed by lattice models”. In: *Physical review letters* 105.21 (2010), p. 218101.

[25] Pavel I Zhuravlev et al. “Propensity to form amyloid fibrils is encoded as excitations in the free energy landscape of monomeric proteins”. In: *Journal of molecular biology* 426.14 (2014), pp. 2653–2666.

[26] D Thirumalai, DK Klimov, and RI Dima. “Emerging ideas on the molecular basis of protein and peptide aggregation”. In: *Current opinion in structural biology* 13.2 (2003), pp. 146–159.

[27] Philipp Neudecker et al. “Structure of an intermediate state in protein folding and aggregation”. In: *Science* 336.6079 (2012), pp. 362–366.

[28] Josh Abramson et al. “Accurate structure prediction of biomolecular interactions with AlphaFold 3”. In: *Nature* (2024), pp. 1–3.

[29] Dylan T Murray et al. “Structure of FUS protein fibrils and its relevance to self-assembly and phase separation of low-complexity domains”. In: *Cell* 171.3 (2017), pp. 615–627.

[30] William Humphrey, Andrew Dalke, and Klaus Schulten. “VMD: visual molecular dynamics”. In: *Journal of molecular graphics* 14.1 (1996), pp. 33–38.

[31] John Jumper et al. “Highly accurate protein structure prediction with AlphaFold”. In: *Nature* 596.7873 (2021), pp. 583–589.

[32] Myungwoon Lee et al. “Molecular structure and interactions within amyloid-like fibrils formed by a low-complexity protein sequence from FUS”. In: *Nature communications* 11.1 (2020), p. 5735.

[33] Laura Esteban-Hofer et al. “Ensemble structure of the N-terminal domain (1–267) of FUS in a biomolecular condensate”. In: *Biophysical Journal* 123.5 (2024), pp. 538–554.

[34] Ashish Joshi et al. “Single-molecule FRET unmasks structural subpopulations and crucial molecular events during FUS low-complexity domain phase separation”. In: *Nature Communications* 14.1 (2023), p. 7331.

Table S2: **Kinetic rate constants.** Reverse rate constants from fibril states $i$ to RC ensemble relative to core 3 (S-bend), computed using MFPTs and relative free energies.

| Fibril State | $\tau_i$ (s) | $\Delta F_{i3}$ (kcal/mol) | $\frac{k_{\text{rev},i}}{k_{\text{rev},3}}$ |
|---|---|---|---|
| 1 | $1.554 \pm 0.016$ | $-2.70 \pm 0.10$ | $(2.79 \pm 0.47) \times 10^{-3}$ |
| 2 | $0.685 \pm 0.010$ | $-1.78 \pm 0.11$ | $(2.99 \pm 0.56) \times 10^{-2}$ |
| 3 | $0.4049 \pm 0.007$ | 0 | 1 (reference) |

Table S3: **AlphaFold quality metrics.** Values of the quality metrics for AlphaFold predictions of $FUS_{155\text{-}190}$ and $FUS_{141\text{-}214}$ fibrils.

| Structure | Average pLDDT | ipTM | pTM |
|---|---|---|---|
| $FUS_{155\text{-}190}$, AF3 | 78.73 | 0.50 | 0.51 |
| $FUS_{141\text{-}214}$, AF3 | 70.71 | 0.55 | 0.55 |
| $FUS_{155\text{-}190}$, AF2 | 23.90 | 0.16 | 0.18 |
| $FUS_{141\text{-}214}$, AF2 | – | – | – |

Table S4: **Distance measurements on sections of FUS-LC.** Distances between selected residues within the same FUS-LC chain, measured in both the dilute and condensed phases. Averages and standard deviations obtained the simulations are compared experimental values [33], measured using double electron–electron resonance (DEER). All values are in nm.

| Residues | Source | Dilute phase | Condensed phase |
|---|---|---|---|
| A10 S29 | Simulations | $2.82 \pm 0.85$ | $2.83 \pm 0.85$ |
| | Experiment | $3.21 \pm 1.04$ | $2.82 \pm 0.83$ |
| S29 S61 | Simulations | $3.67 \pm 1.21$ | $3.73 \pm 1.21$ |
| | Experiment | $4.00 \pm 1.39$ | $3.83 \pm 1.21$ |
| S61 S86 | Simulations | $3.17 \pm 1.01$ | $3.21 \pm 1.01$ |
| | Experiment | $3.52 \pm 1.13$ | $3.19 \pm 1.01$ |
| S86 A105 | Simulations | $2.83 \pm 0.85$ | $2.84 \pm 0.86$ |
| | Experiment | $3.13 \pm 1.00$ | $3.00 \pm 0.88$ |
| A105 G128 | Simulations | $3.03 \pm 0.92$ | $3.06 \pm 0.93$ |
| | Experiment | $3.60 \pm 1.12$ | $3.34 \pm 1.01$ |
| G128 Q158 | Simulations | $3.51 \pm 1.12$ | $3.57 \pm 1.13$ |
| | Experiment | $4.42 \pm 1.47$ | $3.80 \pm 1.15$ |
| Q158 M184 | Simulations | $3.14 \pm 1.05$ | $3.17 \pm 1.05$ |
| | Experiment | $3.66 \pm 1.20$ | $3.54 \pm 1.05$ |
| M184 S205 | Simulations | $2.81 \pm 0.94$ | $2.84 \pm 0.95$ |
| | Experiment | $3.31 \pm 1.04$ | $3.13 \pm 0.89$ |

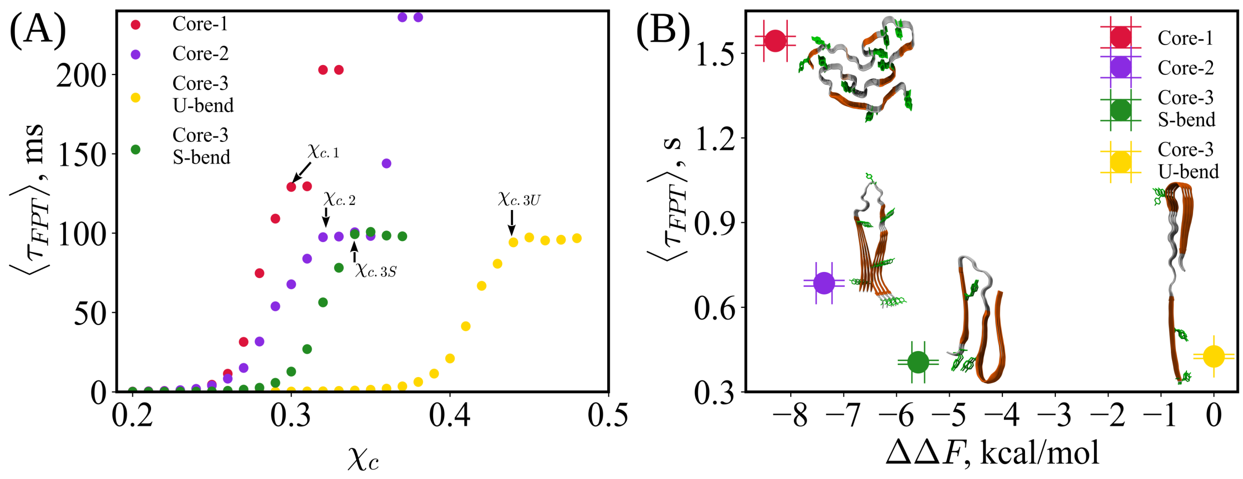


Figure S1: **Structural overlap parameter ($\chi_{fib}$) distributions and their relation to first passage times in FUS variants.** (A) Mean first passage times $\langle \tau_{FPT} \rangle$ as a function of the structural overlap cutoff $\chi_c$ for different cores. (B) MFPTs as a function of relative stability with respect to core-3 (U-bend) for FUS-LC. The inset shows the experimental structures of core-1 (PDB ID: 5W3N [29]) and core-2 (PDB ID: 6XFM [32]), along with the predicted S-bend and U-bend structures of core-3.

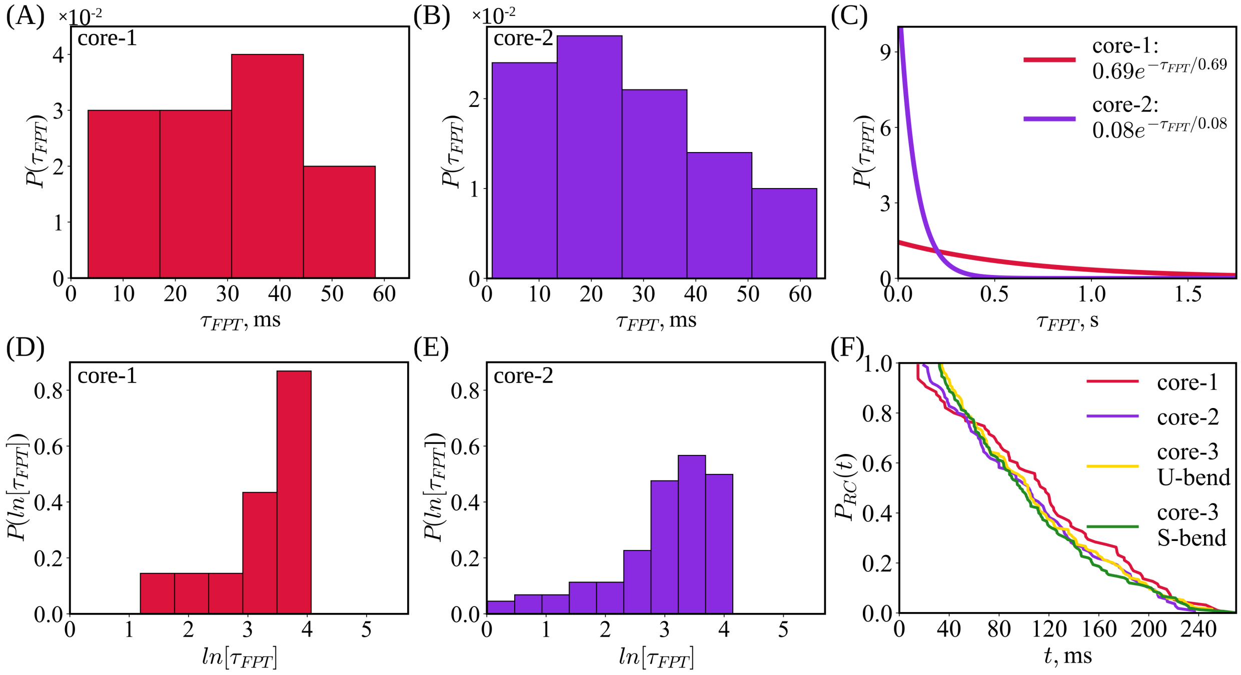


Figure S2: **Ostwald rule of stages (extended).** (A-B) Distributions of first-passage times (FPTs) for the transition from the disordered ground state to the fibril-like $N^*$ states in FUS-LC-N (residues 1–163). Shown are the $\tau_{FPT}$ distributions sampled from Brownian dynamics simulations (see Methods) for the transition from the free energy ground state of FUS-LC to the core-1 fibril-like structure (panel A), and to the core-2 fibril-like structure (panel B). (C) Theoretical FPT distributions for FUS-LC-N computed using Eq. 2 in the main text. (D–E) Distributions of $ln[\tau_{FPT}]$ for RC $\rightarrow$ $N^*$ transitions in FUS-LC-N. (F) Probability of FUS-LC being in a random coil conformation at time $t$.

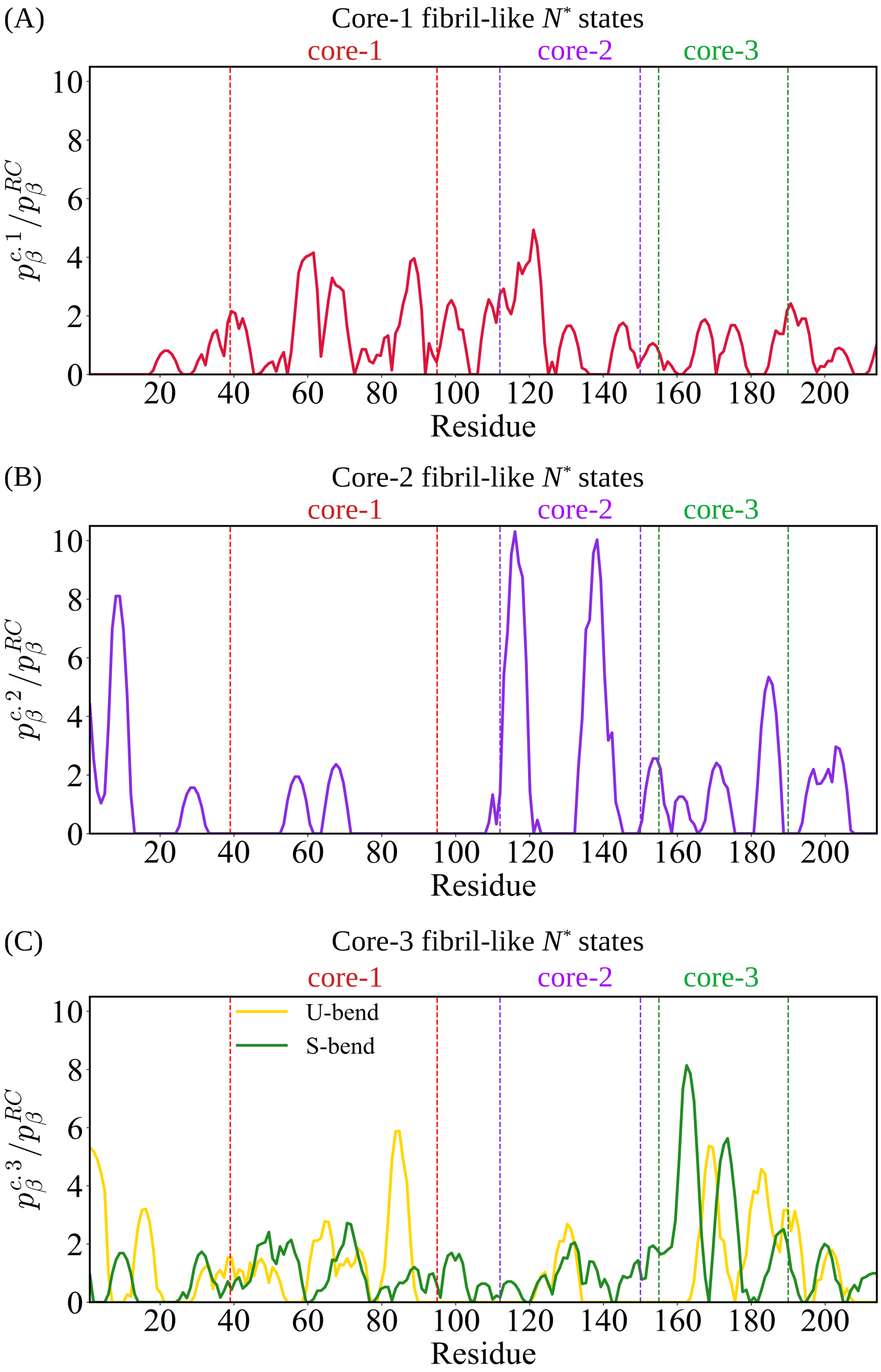

(A)
Core-1 fibril-like $N^*$ states
core-1
core-2
core-3
$p_\beta^{c.1}/p_\beta^{RC}$
Residue
(B)
Core-2 fibril-like $N^*$ states
core-1
core-2
core-3
$p_\beta^{c.2}/p_\beta^{RC}$
Residue
(C)
Core-3 fibril-like $N^*$ states
core-1
core-2
core-3
U-bend
S-bend
$p_\beta^{c.3}/p_\beta^{RC}$
Residue

Figure S3: **Residue-wise enhancement of $\beta$-strand propensity in fibril-like $N^*$ states.** Ensemble-averaged, residue-wise ratios of the propensity to form $\beta$ strand in the $N^*$ states, $p_\beta^{c,i}$ ($i = 1, 2, 3$), to the corresponding values in the remaining conformations of the ensemble, $p_\beta^{RC}$. (A) Ratio for core-1 fibril-like $N^*$ states, panel (B) for core-2, and panel (C) for core-3, including both S-bend and U-bend fibril-like conformations. Vertical dashed lines indicate the residue boundaries of each core region: core-1 (residues 39–95), core-2 (residues 112–150), and core-3 (residues 155–190).

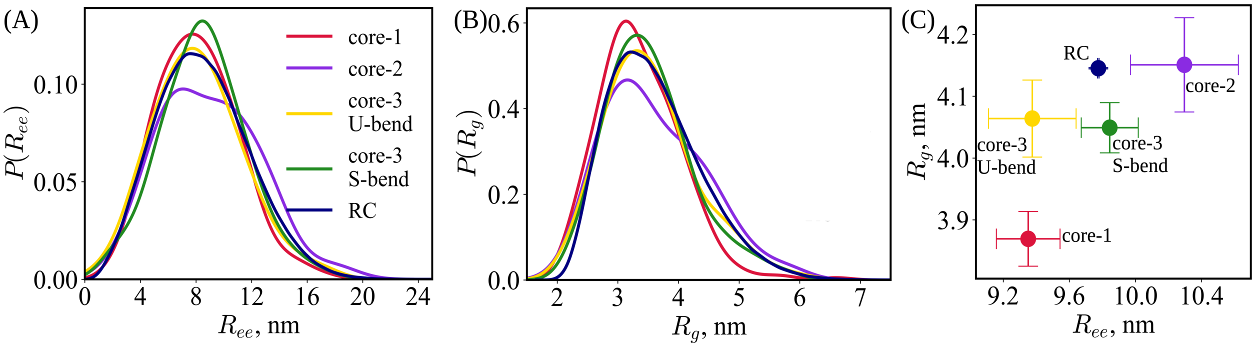


Figure S4: **Conformations of monomers in fibril-like $N^*$ states.** Probability density distributions of the end-to-end distance, $R_{ee}$ (panel A), and the radius of gyration, $R_g$ (panel B), for FUS-LC monomers. Distributions are computed separately for fibril-like $N^*$ conformations ($\chi_i \geq \chi_c$) and for random coil (RC) conformations ($\chi_{fib.i} < \chi_c$). Panel (C) shows the average values of $R_{ee}$ and $R_g$ for each of these states, highlighting structural differences between fibril-like and random-coil ensembles.

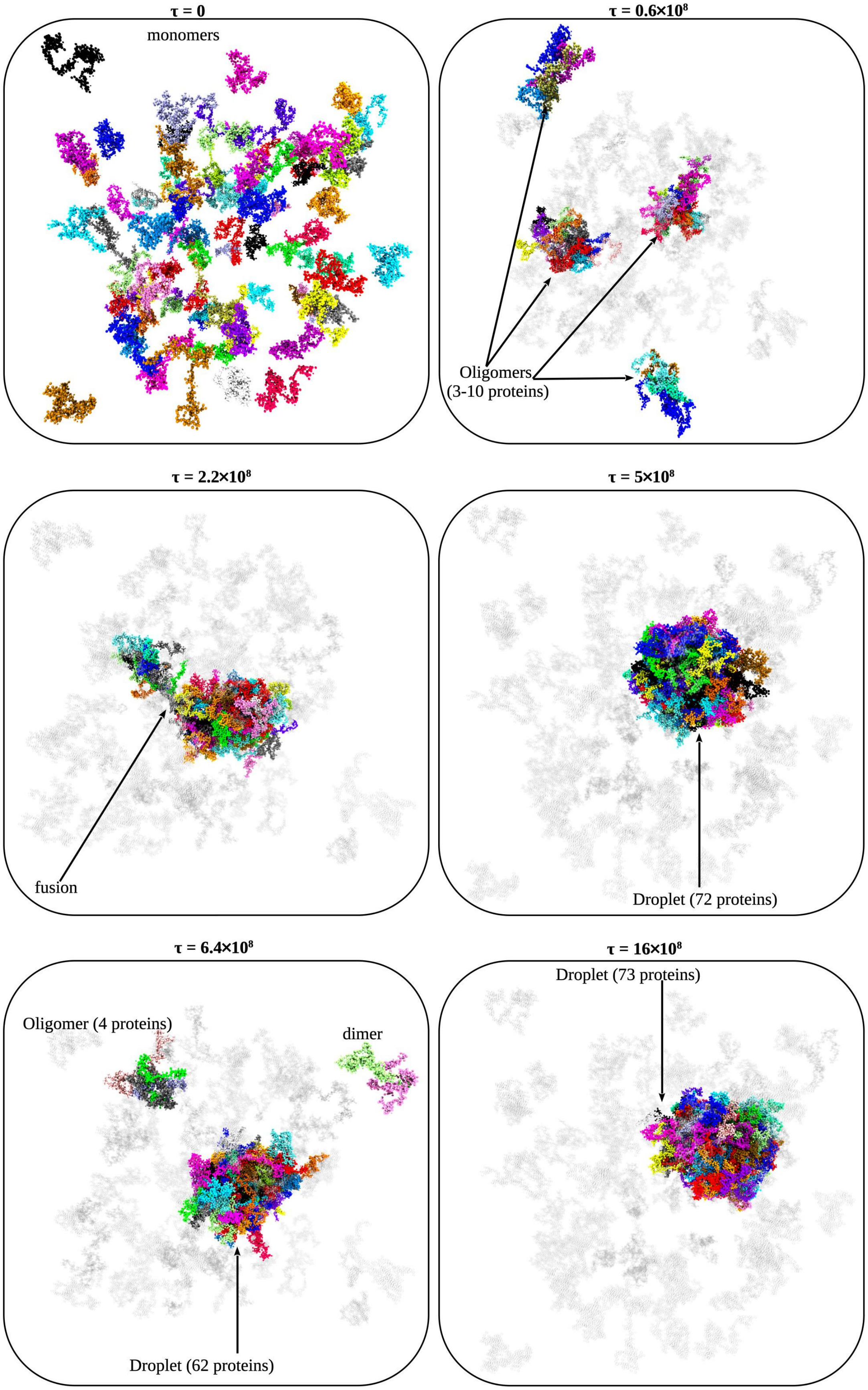


Figure S5: **Phase separation in FUS-LC.** Images from Langevin simulations of 96 FUS-LC chains at selected times $\tau$ (see Results in the main text).

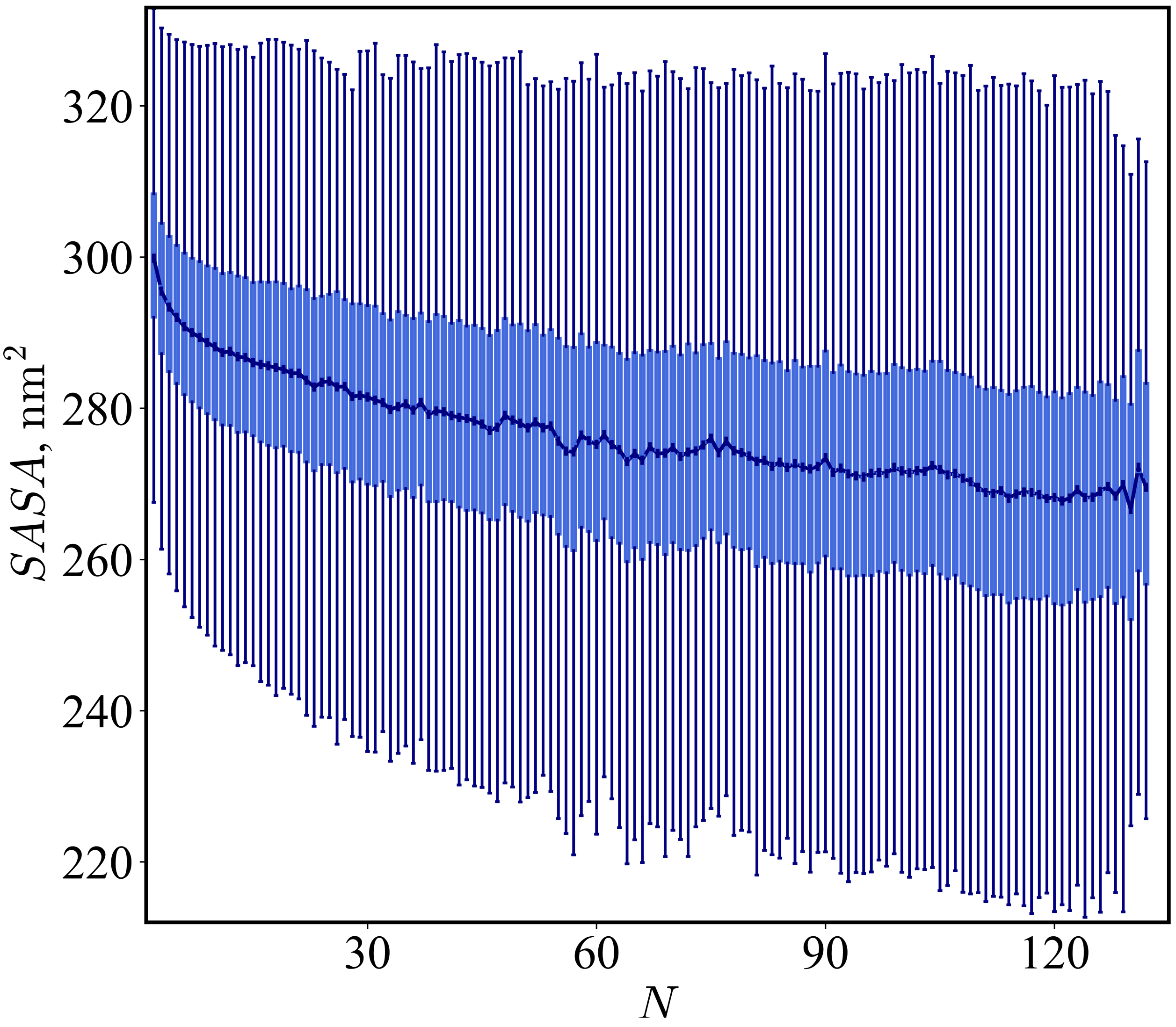


Figure S6: **Monomer Solvent Accessibility as a Function of Assembly Size.** For each assembly size $N$—ranging from dimers to large droplets ($N > 70$)—boxplots show the distribution of solvent-accessible surface area ($SASA$) for a FUS-LC monomer within the corresponding assembly.

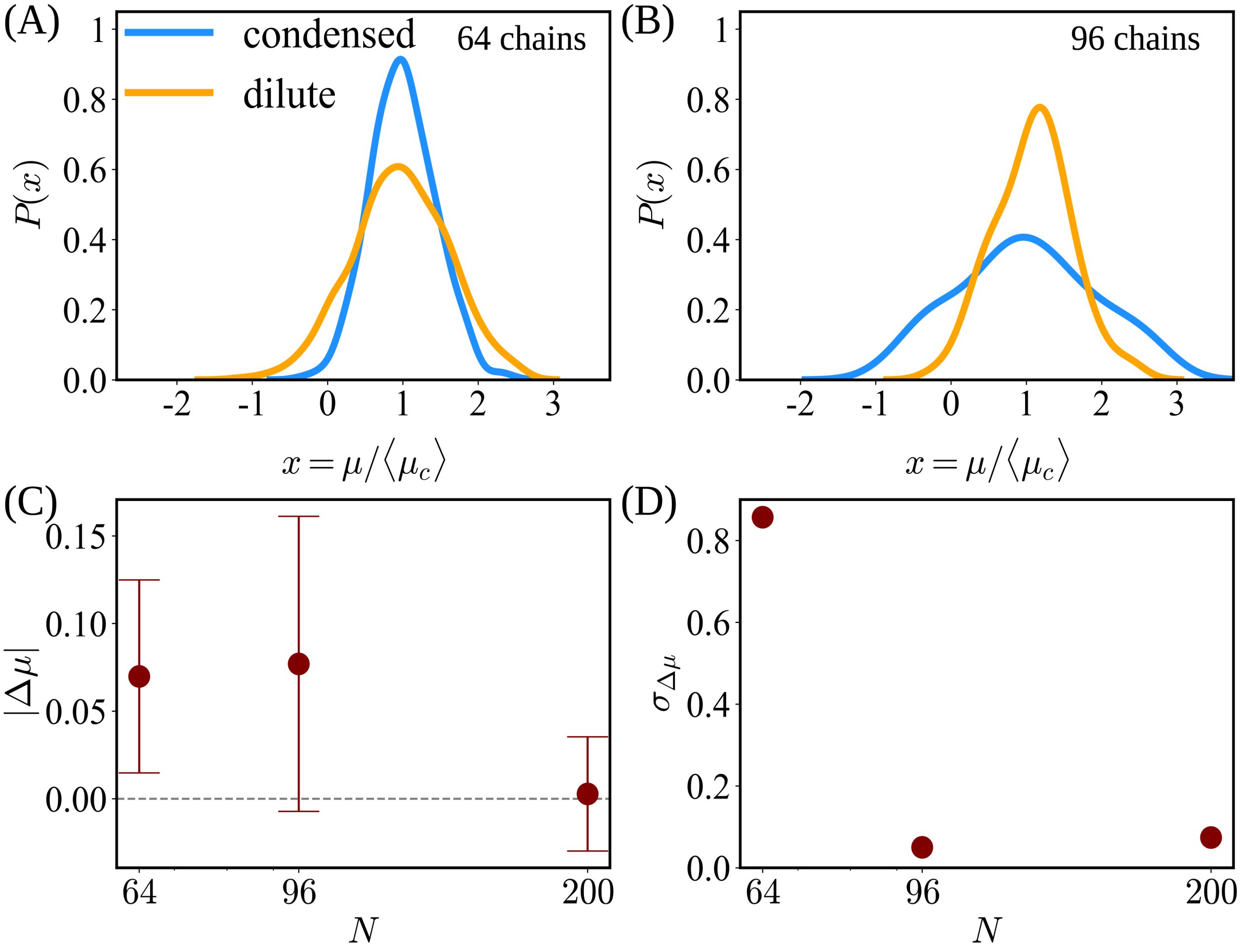


Figure S7: **Distributions of chemical potentials.** Distributions of the chemical potential in the dispersed and condensed phases, scaled by the average chemical potential in the condensed phase, $\langle \mu_c \rangle$, for simulations of 64 (panel A and 96 (panel B) FUS-LC chains. The corresponding distribution for a system of 200 chains is presented in Fig. 3 of the main text. (C) Differences in chemical potential, $|\Delta\mu| = |\mu_c - \mu_d|/\mu_c$, between the condensed and dilute phases as a function of system size $N$, illustrating finite-size effects. (D) Dependence of $\sigma_{\Delta\mu}$ on $N$(see Eq. 15 in SI Methods).

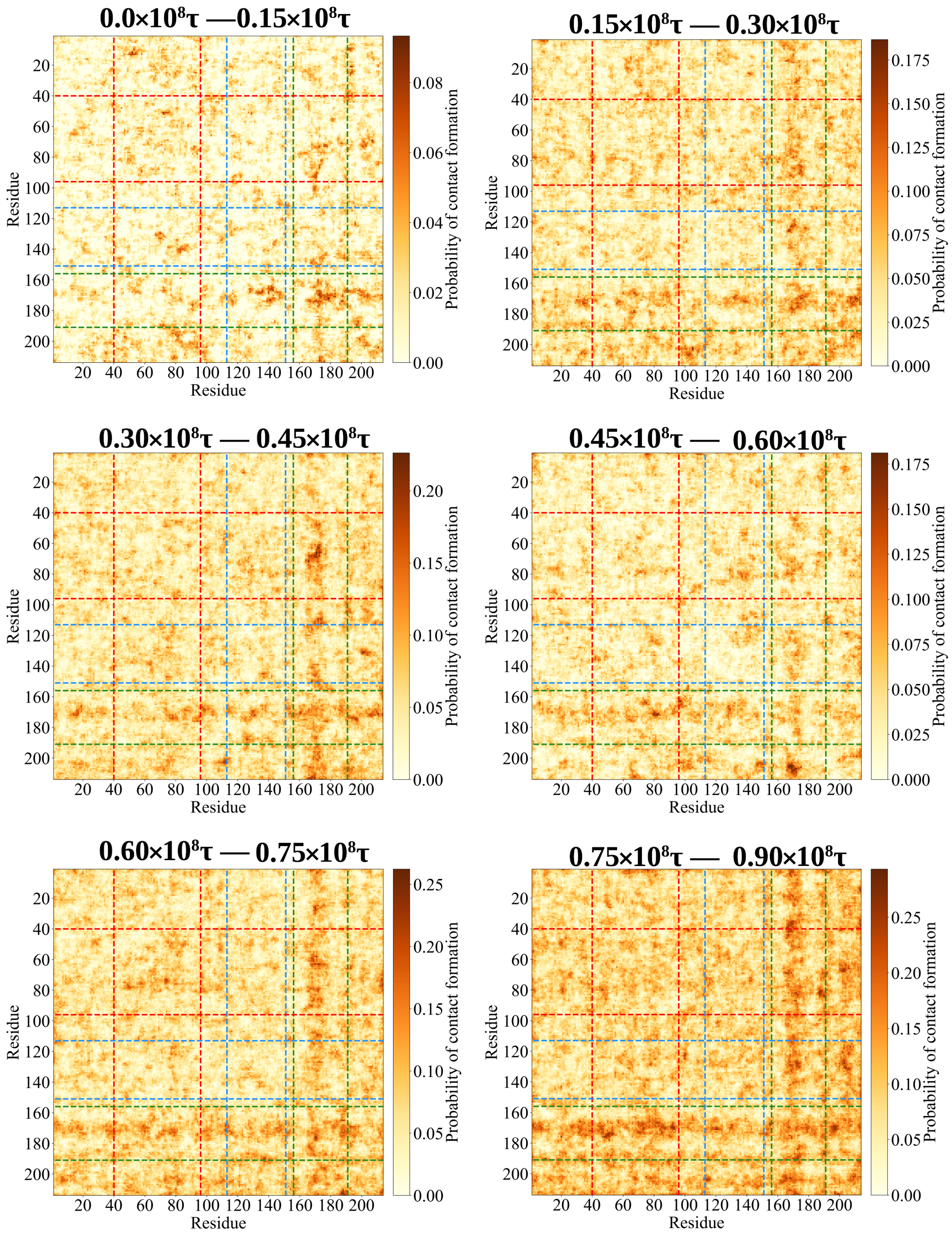


Figure S8: **Evolution of interchain interactions during phase separation.** Contact maps for chains within small oligomers and droplets, averaged over specified simulation time intervals ($\tau$). Dashed lines are the regions of the amino acid sequence corresponding to core-1 (red), core-2 (blue), and core-3 (green).

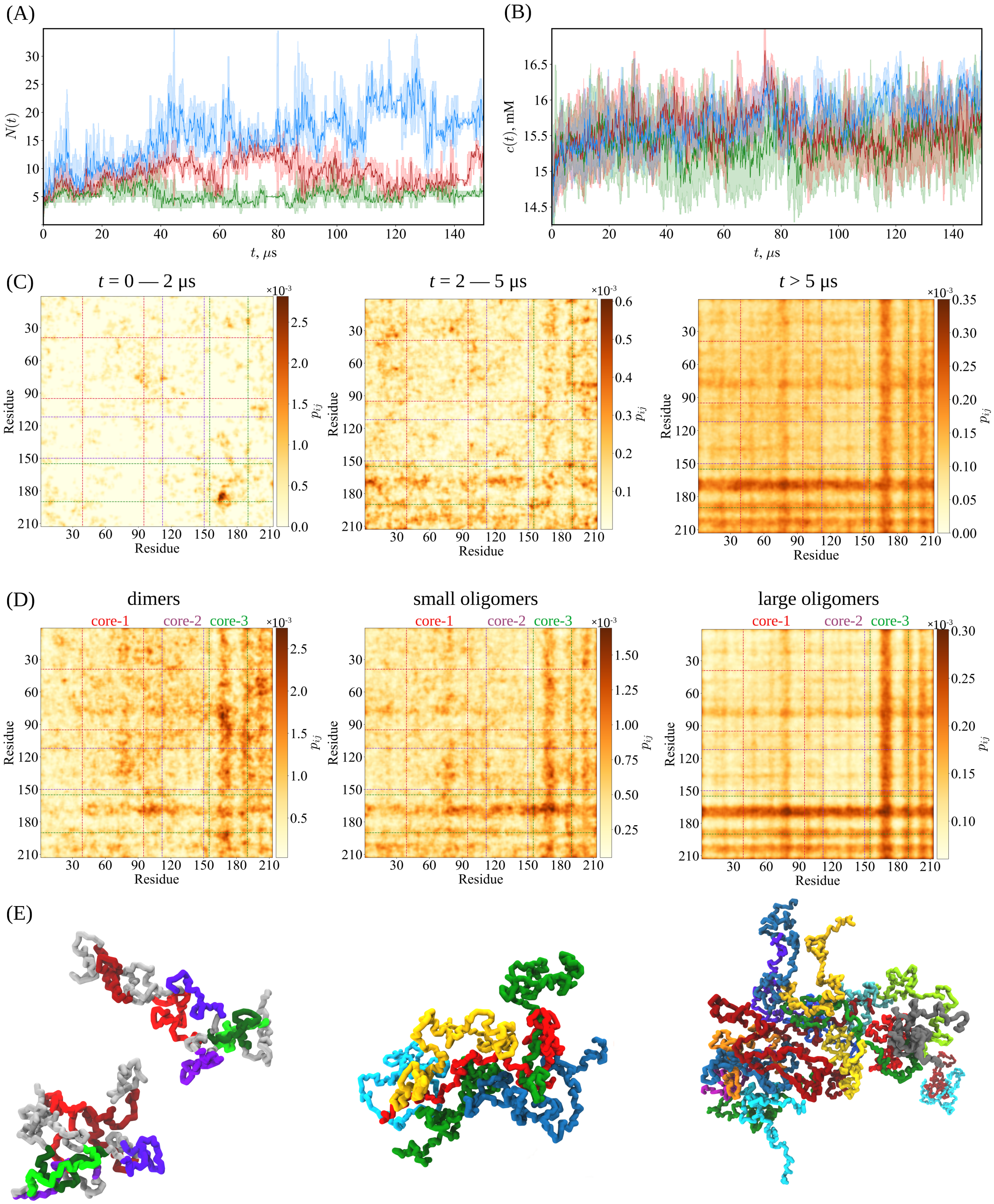

(A)
N(t)
t, μs
(B)
c(t), mM
t, μs
(C)
t = 0 — 2 μs
t = 2 — 5 μs
t > 5 μs
Residue
$p_{ij}$
(D)
dimers
small oligomers
large oligomers
core-1
core-2
core-3
(E)

Figure S9: **Brownian dynamics of Phase separation probed by Brownian dynamics .** Results from 150 $\mu$s long Brownian dynamics simulation consisting of 64 FUS-LC proteins during phase separation. (A) Temporal evolution of the sizes of the three largest oligomers (blue, red, and green) corresponding to the trajectories in panel B. Droplet size is defined as the number of FUS-LC chains contained within each oligomer. (B) Protein mass concentration $c$ as a function of simulation time for the oligomer shown in panel A. (C) Contact maps for chains within small oligomers and droplets, averaged over selected simulation time intervals $t$. Dashed lines are the regions of the amino acid sequence corresponding to core-1 (red), core-2 (violet), and core-3 (green). (D) Contact maps illustrating inter-chain interactions: dimers (left), small oligomers ($N < 10$ chains; middle), and large oligomers ($10 \leq N \leq 31$ chains; right). (E) Representative conformations of FUS-LC in assemblies corresponding to those in panel (D). The left panel shows two dimers, where monomers interact primarily via core-3 regions (green). Core-1 and core-2 regions are highlighted in red and violet, respectively, while all other residues are shown in gray.

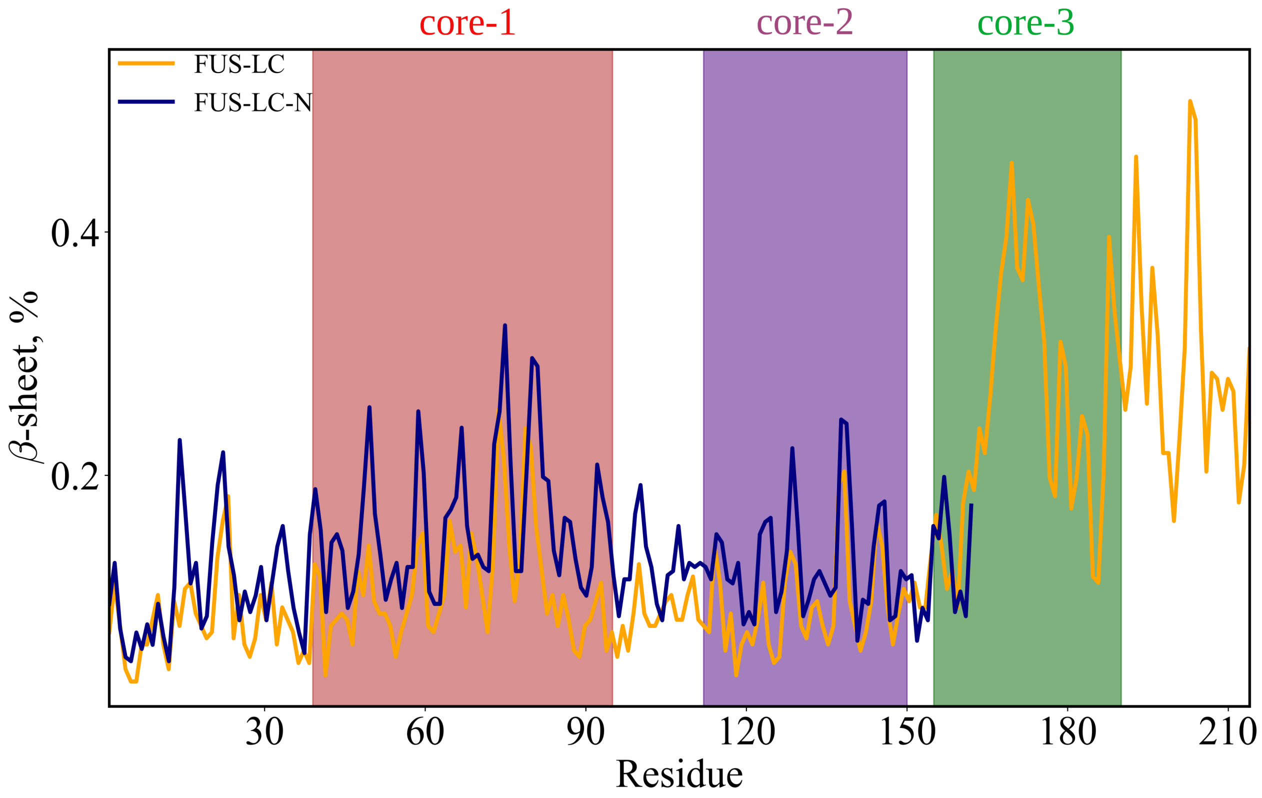


Figure S10: **$\beta$-strand content in the conformational ensembles of FUS monomers.** Residue-wise ensemble-averaged percentages for forming $\beta$-strand for FUS-LC (residues 1-214), FUS-LC-N (residues 1-163), and FUS-LC-C (residues 111-214). Vertical dashed lines are positioned at the residues corresponding to the boundaries of the core regions: core-1 (residues 39-95), core-2 (residues 112-150), and core-3 (residues 155-190).

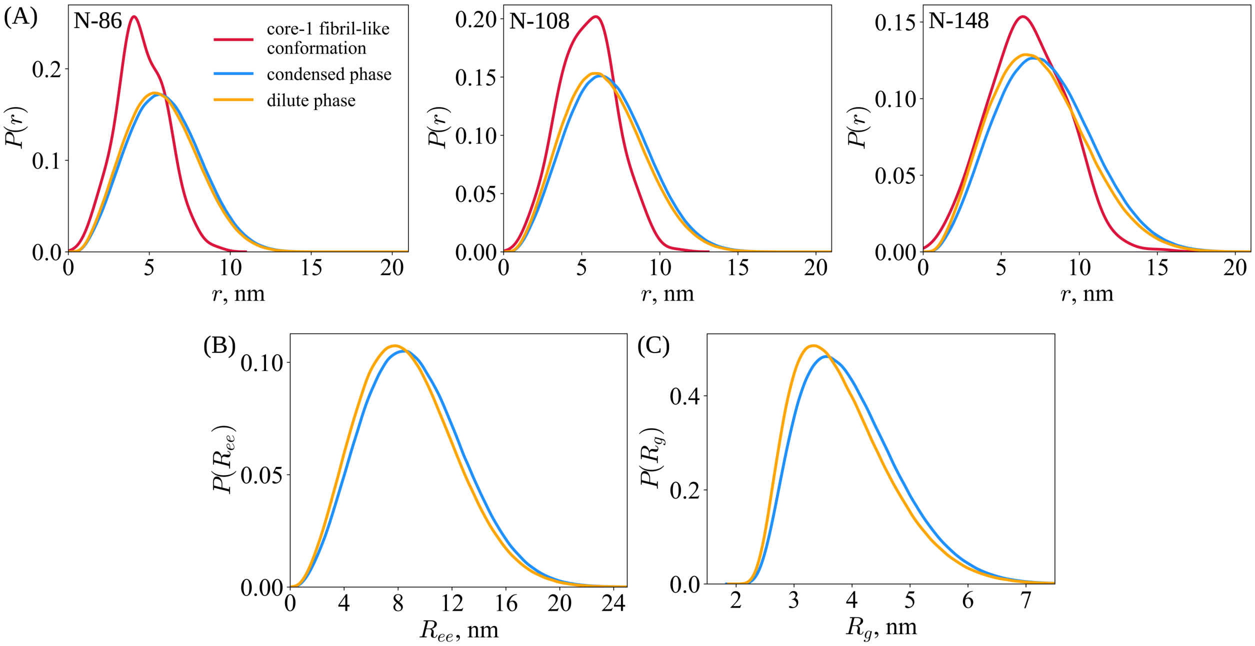


Figure S11: **Conformations of monomers in dispersed and dense phases are similar.** (A) Distributions of distances between selected residue pairs within the same FUS-LC chain, measured separately for chains in the dilute phase and those within the condensed droplet. The residue pairs correspond to the labeled positions in smFRET experiments [34]. The same distances were also separately calculated in core-1 fibril-like conformations. (B-C) Probability density distributions of the radius of gyration ($R_g$) and end-to-end distance ($R_{ee}$) for FUS-LC monomers. These conformational parameters were computed individually for each chain in both the condensed and dilute phases, as well as for monomers in core-1 fibril-like conformations.

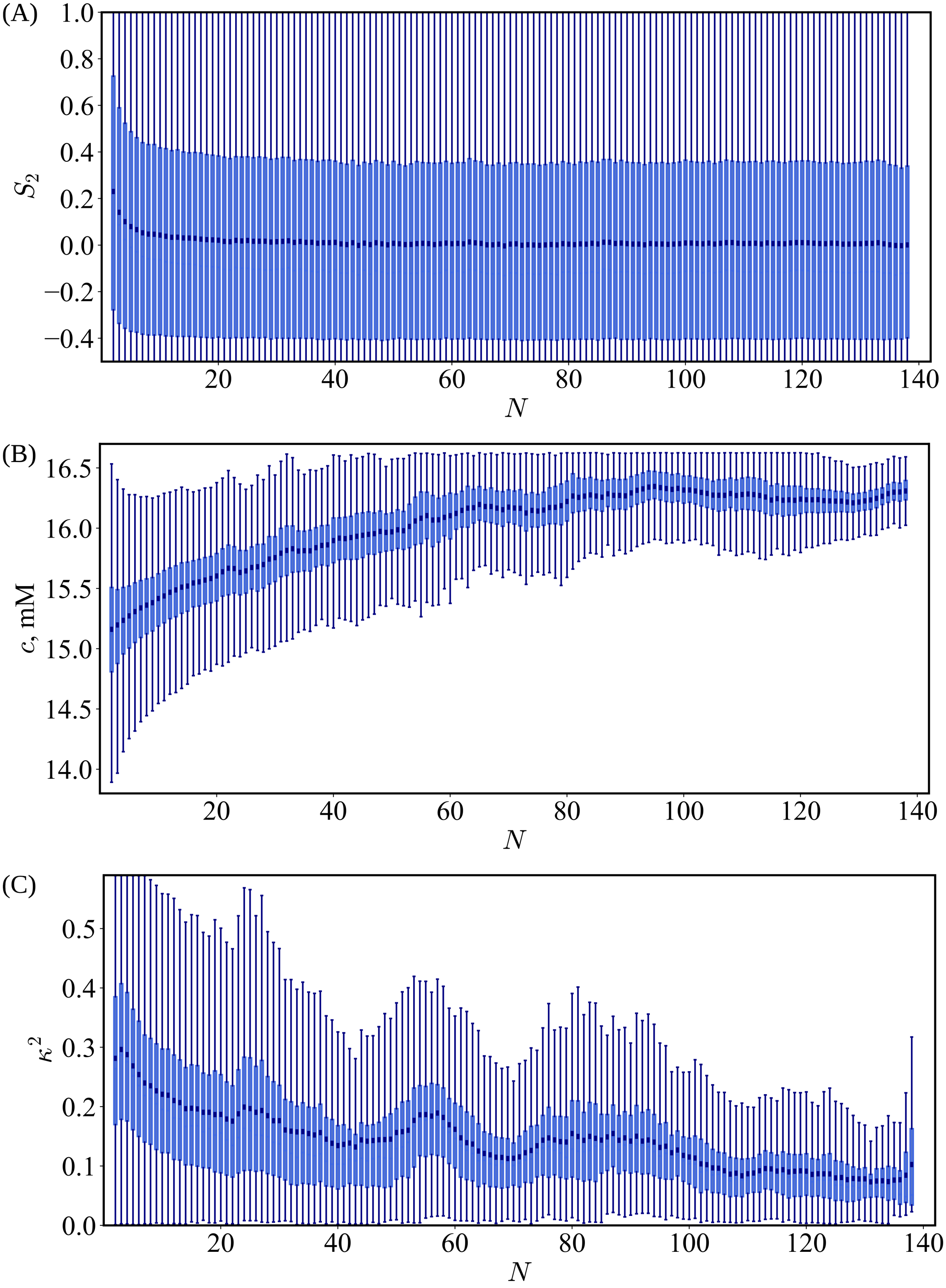
(A)
$S_2$
$N$
(B)
$c$, mM
$N$
(C)
$\kappa^2$
$N$

Figure S12: **Characterization of FUS-LC protein assemblies as a function of assembly size.** For each assembly size $N$—ranging from dimers to droplets ($N > 70$)—boxplots show the distributions of structural properties of FUS-LC assemblies: nematic order parameter $S_2$ (panel A), protein mass concentration within the assembly $c$ (panel B), relative shape anisotropy $\kappa^2$ (panel C), and the shape parameter $S$ (panel D).

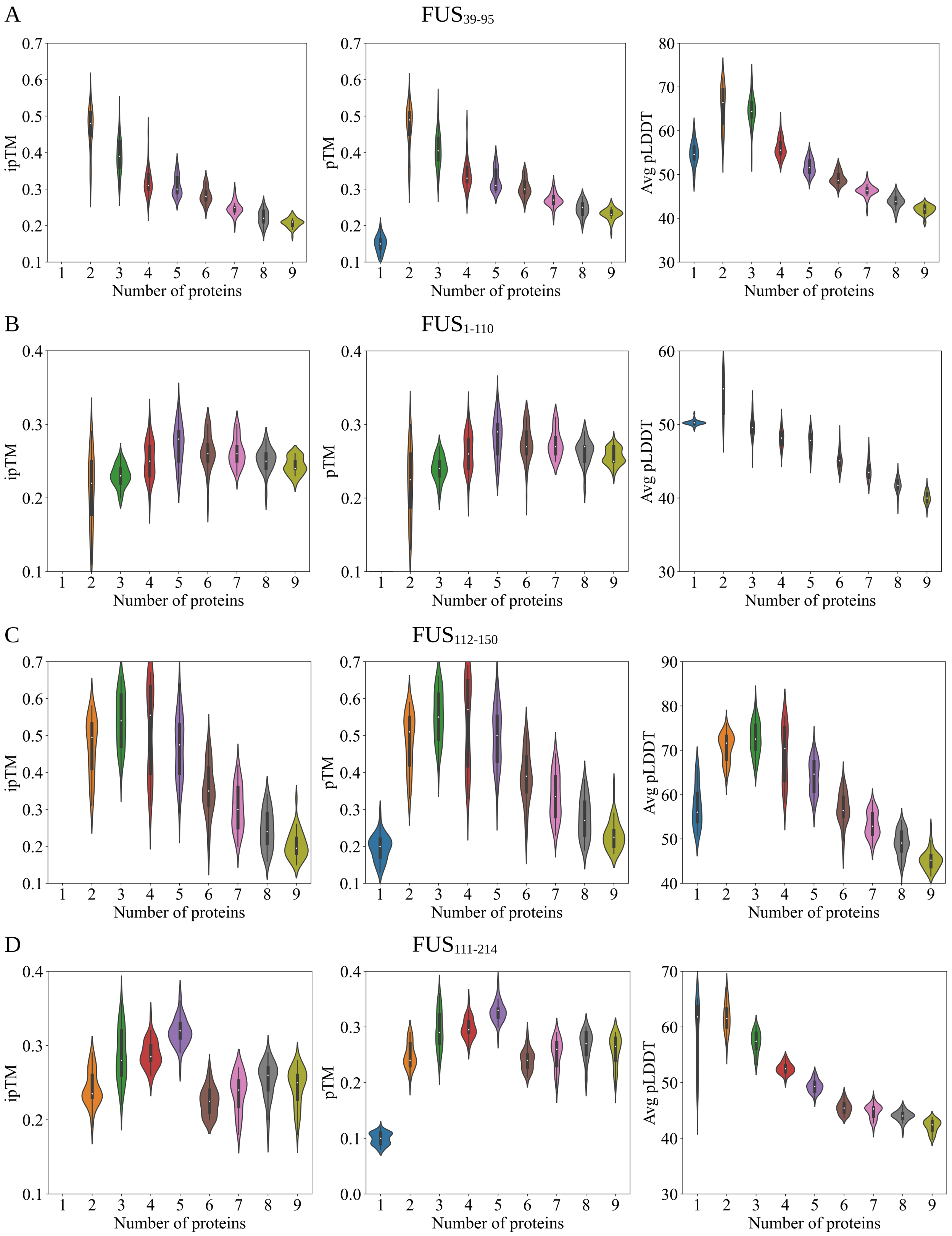
A
FUS39-95
ipTM
pTM
Avg pLDDT
Number of proteins
B
FUS1-110
ipTM
pTM
Avg pLDDT
Number of proteins
C
FUS112-150
ipTM
pTM
Avg pLDDT
Number of proteins
D
FUS111-214
ipTM
pTM
Avg pLDDT
Number of proteins

Figure S13: **Distributions of quality metrics for AlphaFold3 predictions of core-1 and core-2 fibrils.** Violin plots display the distributions of Interface predicted template modeling (TM) score (iPTM), predicted TM-score (pTM), and predicted local distance difference test (pLDDT) metrics calculated by AlphaFold server for fibrils formed by varying numbers of FUS peptides. Each metric reflects the confidence in predicted inter- and intra-chain interactions within fibrillar assemblies.Distributions of structural metrics for different FUS fibril constructs: (A) $FUS_{39–95}$, (B) $FUS_{1–110}$, (C) $FUS_{111–214}$, and (D) $FUS_{112–150}$.

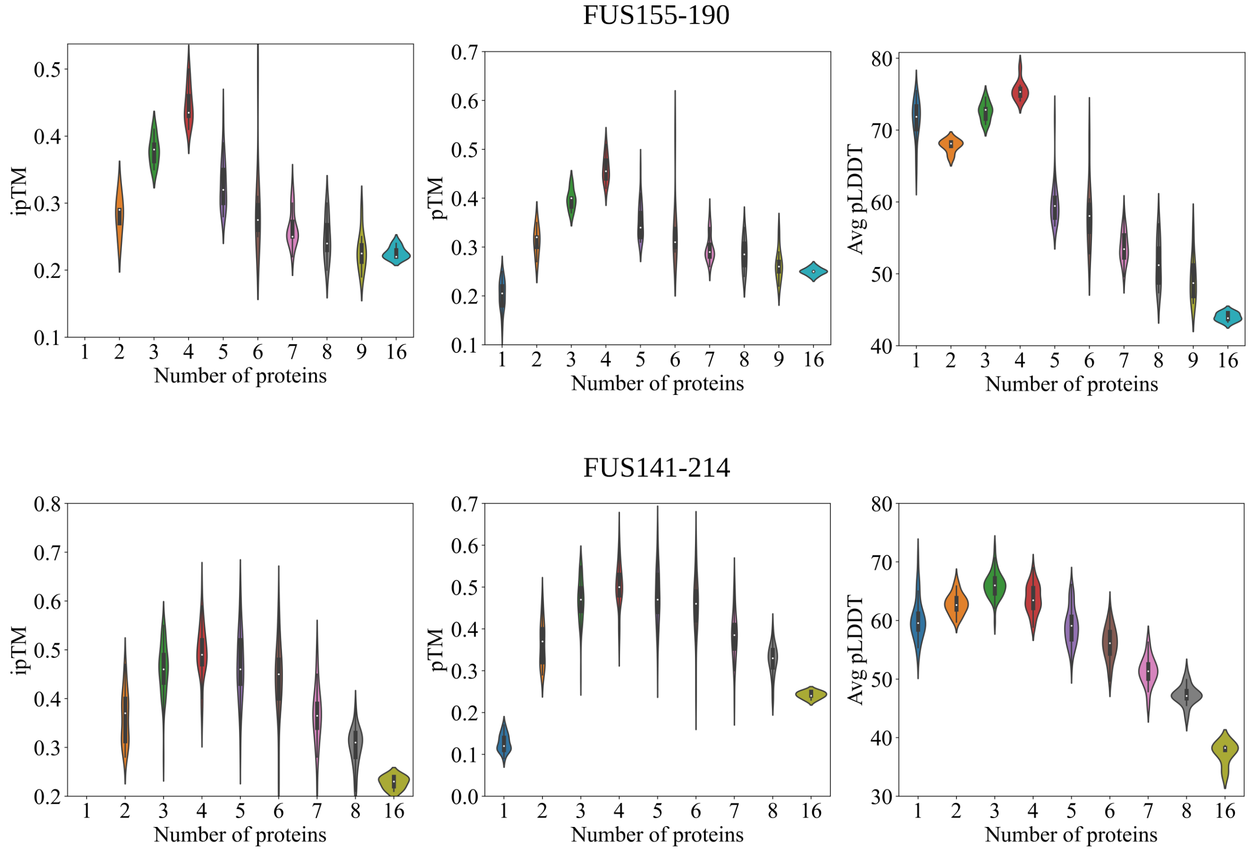


Figure S14: **Distributions of quality metrics for AlphaFold3 predictions of core-3 fibrils.** Same as Fig. S13 but for $FUS_{155\text{-}190}$ and $FUS_{141\text{-}214}$ fibrils.

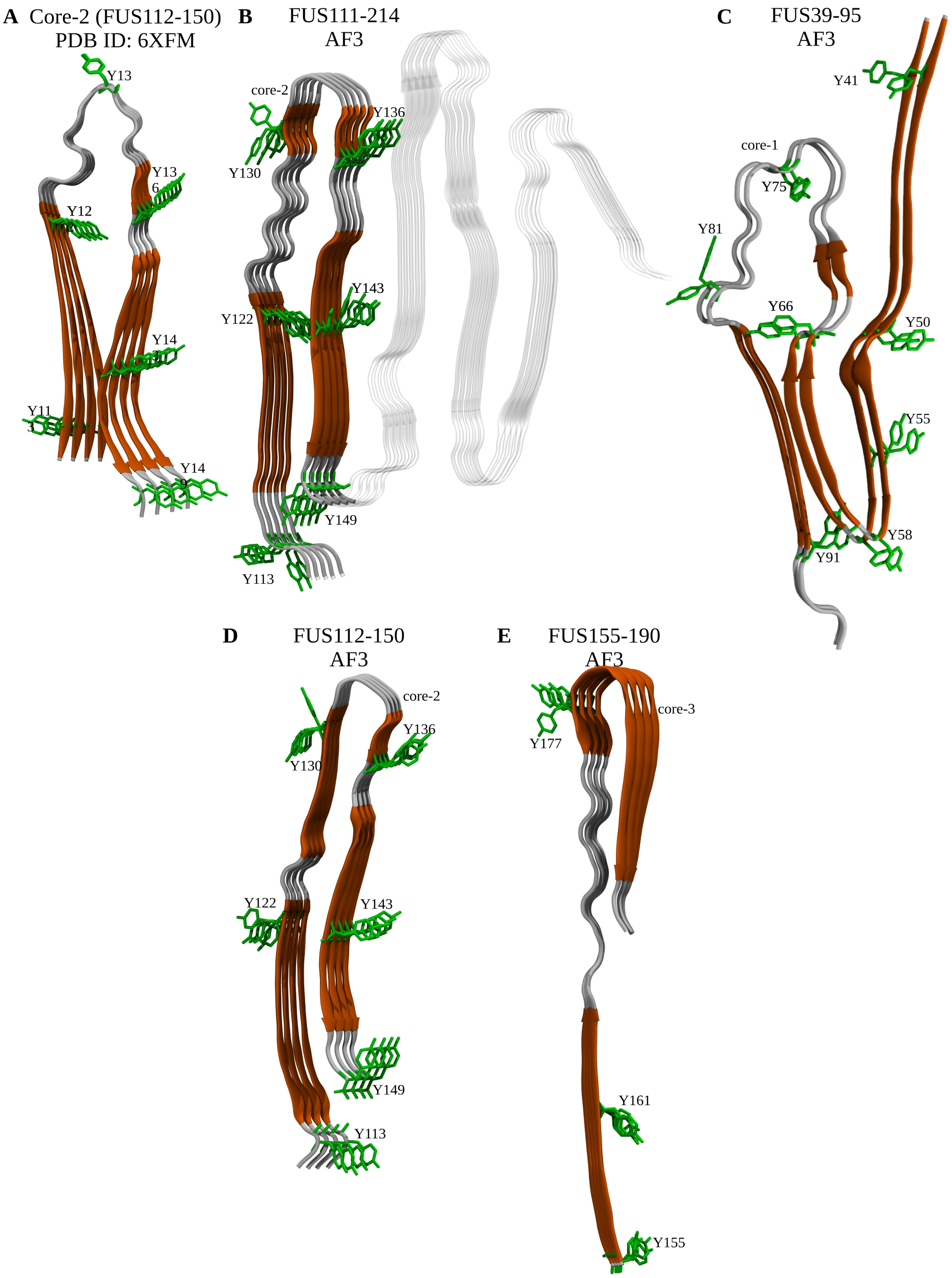

A Core-2 (FUS112-150)
PDB ID: 6XFM
Y13
Y13
6
Y12
Y14
Y11
Y14
B FUS111-214
AF3
core-2
Y136
Y130
Y143
Y122
Y149
Y113
C FUS39-95
AF3
Y41
core-1
Y75
Y81
Y66
Y50
Y55
Y58
Y91
D FUS112-150
AF3
core-2
Y136
Y130
Y122
Y143
Y149
Y113
E FUS155-190
AF3
core-3
Y177
Y161
Y155

**Figure S15: Structural models of core-2 and core-3 fibrils predicted by AlphaFold3.** (A) Structure of core-2 (residues 112–150) resolved by cryo-EM [32]. (B, C) Predicted pentameric (B) and tetrameric (C) assemblies of $FUS_{111\text{-}214}$ and $FUS_{112\text{-}150}$, respectively. In panel (B), core-2 residues (112–150) are depicted as in panel (A), while the remainder of the fibril is shown in transparent for clarity. (D) Structure of a tetrameric assembly of $FUS_{155\text{-}190}$ chains as predicted by AlphaFold3. The color scheme in all panels is the same as in Fig. 5 in the main text.

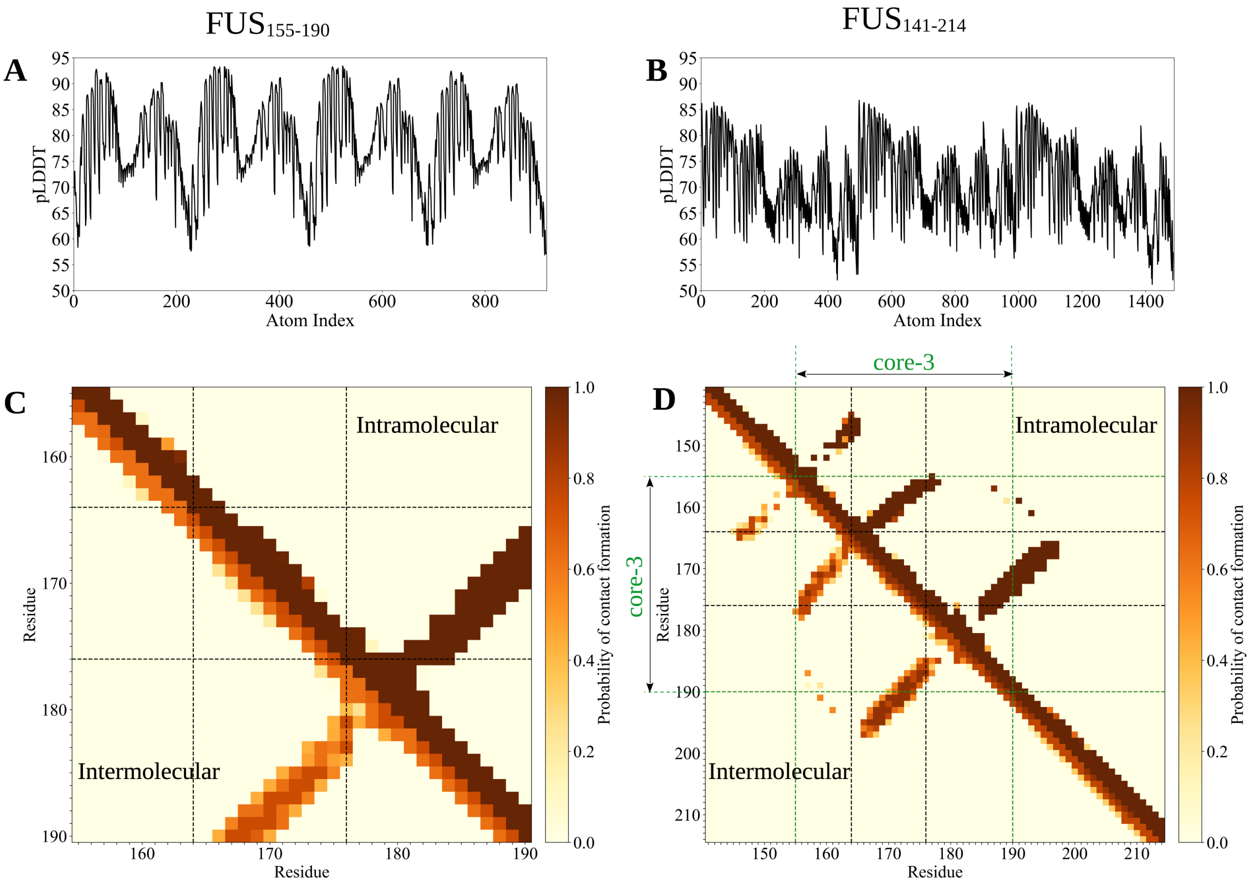


Figure S16: **Assessment of AlphaFold3 predictions of FUS$_{155\text{-}190}$ and FUS$_{141\text{-}214}$ fibrils.** (A) and (B) Predicted local distance difference test (pLDDT) scores calculated by the AlphaFold server for the FUS$_{155\text{-}190}$ (Fig. S15E) and FUS$_{141\text{-}214}$ fibrils shown in Fig. 5C, reflecting confidence in structural predictions. (C) and (D) Contact maps for the fibrils in panels (A) and (B), respectively, where intermolecular contacts are shown in the lower triangle and intramolecular contacts in the upper triangle. Dashed lines highlight the amino acid residues 164-176, where monomer simulations indicated the highest propensity for $\beta$-strand formation (see Fig. S10 in the main text) and droplet simulations showed the greatest probability of interchain contact formation (see Fig. 3 in the main text).

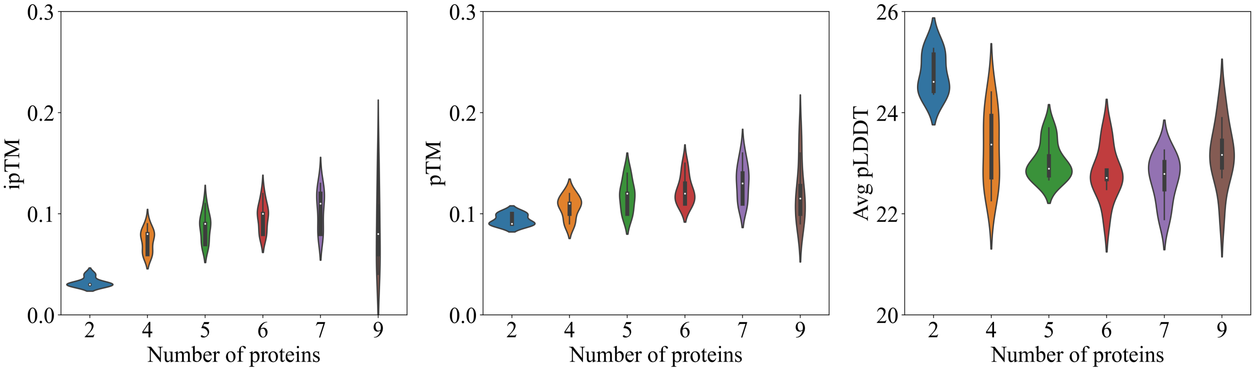


Figure S17: **Distributions of quality metrics for AlphaFold2 predictions of $\mathbf{FUS_{155\text{-}190}}$ fibrils.** Same as in Fig. S14 but for AF2.

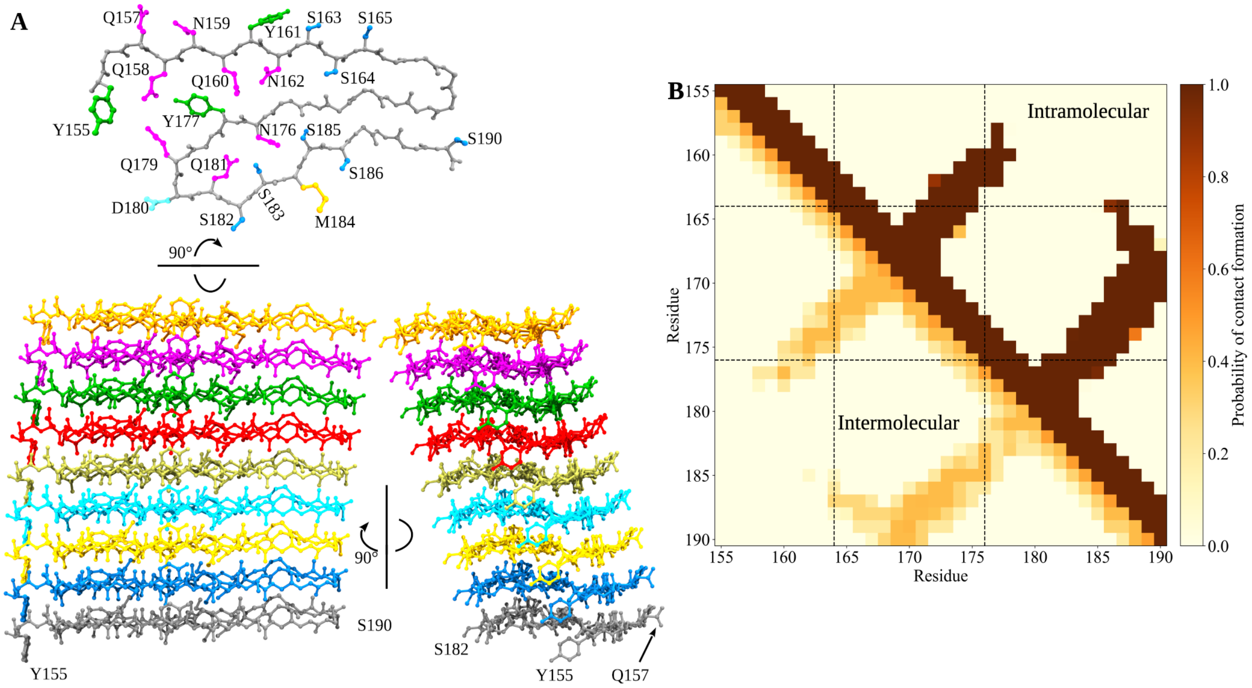


Figure S18: **Structure of CTC predicted by AlphaFold3 for 9 chains of $\mathbf{FUS_{155\text{-}190}}$.** (A) Side (top) and two cross-sectional (bottom) views of the predicted structure model of the complex comprising 9 $FUS_{155\text{-}190}$ peptides predicted using AlphaFold3. The side chains in the top panel of SER, TYR, ASP, and MET residues are colored blue, green, cyan, and yellow, respectively. The side chains of GLN and ASN residues are shown in magenta, with the backbone and GLY residues in grey. (B) Contact maps of the complex shown in panel A: lower and upper triangles represent intermolecular and intramolecular interactions, respectively. Dashed lines highlight the amino acid residues 164-176, where monomer simulations indicated the highest propensity for $\beta$-strand formation (see Fig. S10 in the main text) and droplet simulations showed the greatest probability of interchain contact formation (see Fig. 3 in the main text).

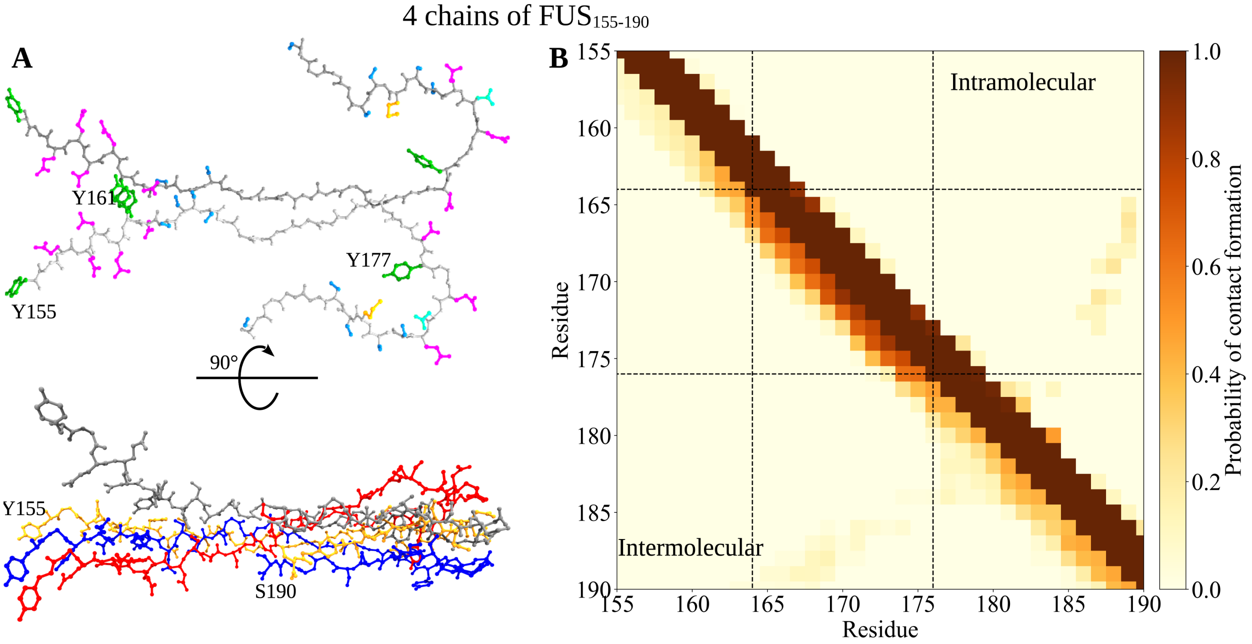


Figure S19: **Structure of core-3 predicted using AlphaFold2.** (A) Cross-sectional (top) and side (bottom) views of the predicted structural model of the $FUS_{155\text{-}190}$ protein complex. The side chains of SER, TYR, ASP, and MET residues are colored blue, green, cyan, and yellow, respectively. The side chains of GLN and ASN residues are depicted in magenta. The backbone and GLY residues are in grey. (B) The contact maps illustrate intermolecular (lower triangle) and intramolecular (upper triangle) interactions. The dashed lines highlight the amino acid residues 164-176, where monomer simulations showed the highest propensity for $\beta$-strand formation (see Fig. S10)and droplet simulations indicated the greatest probability of interchain contact formation (see Fig. 3).

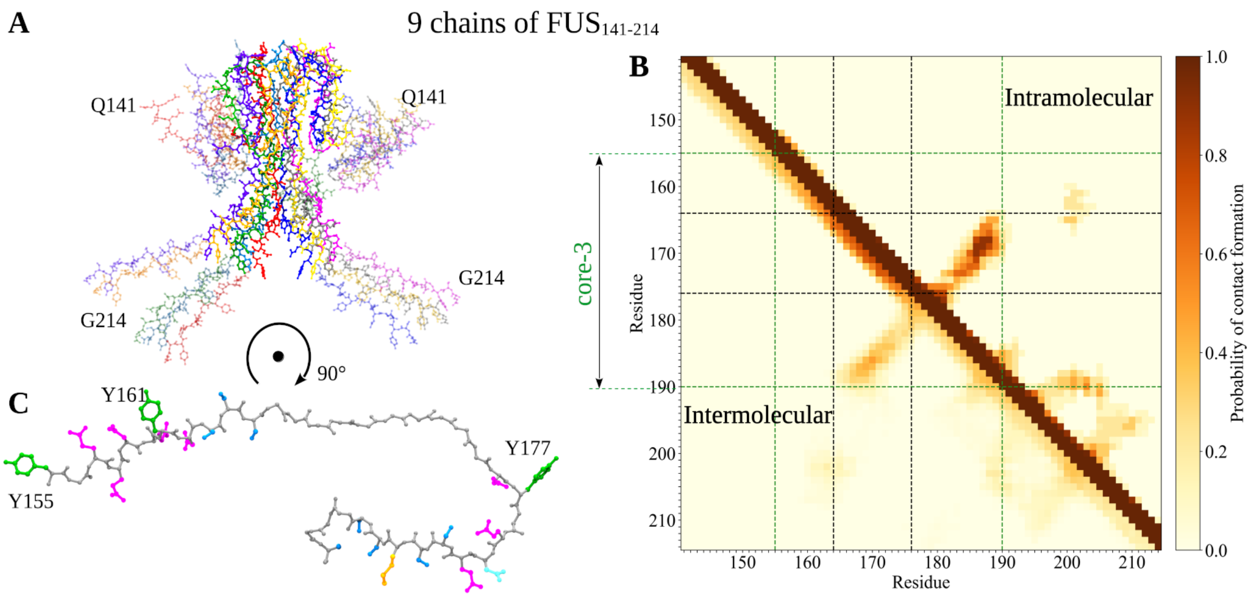


Figure S20: **Structure of $\text{FUS}_{141\text{-}214}$ predicted by AlphaFold2.** (A) Predicted structure of a nine-protein $\text{FUS}_{141\text{-}214}$ complex. Core-3 residues (155-190) are highlighted in opaque representation. (B) Contact map of the complex. Lower and upper triangles represent inter- and intramolecular contacts, respectively. Green dashed lines indicate core-3 boundaries; black dashed lines correspond to those in Fig. S18 in the main text. (C) Side view of core-3 (residues 155 to 190) from one of the monomers $\text{FUS}_{141\text{-}214}$ in panel A with residue coloring as in Fig. S19.

**Supplementary Movie 1: Oligomer Nucleation and Growth.** This movie shows the progressive formation and growth of oligomers from an initially homogeneous, dispersed phase.

**Supplementary Movie 2: Ostwald Ripening.** This movie illustrates the process of Ostwald ripening, where the largest oligomer progressively grows at the expense of smaller ones. The smaller oligomers, which appear and dissolve over time, are highlighted with red dashed lines.

**Supplementary Movie 3: Dynamics of the Droplet.** This movie depicts the equilibrium dynamics of a phase-separated droplet in a system of 200 FUS-LC proteins, characterized by approximately equal chemical potentials in the condensed and dilute phases ($\langle \mu_c \rangle - \langle \mu_d \rangle \approx 0$). The movie captures protein chains entering and exiting the droplet, as well as diffusing within the condensed phase.
